# A new family of structurally conserved fungal effectors displays epistatic interactions with plant resistance proteins

**DOI:** 10.1101/2020.12.17.423041

**Authors:** Noureddine Lazar, Carl H. Mesarich, Yohann Petit-Houdenot, Nacera Talbi, Ines Li de la Sierra-Gallay, Emilie Zélie, Karine Blondeau, Jérôme Gracy, Bénédicte Ollivier, Françoise Blaise, Thierry Rouxel, Marie-Hélène Balesdent, Alexander Idnurm, Herman van Tilbeurgh, Isabelle Fudal

## Abstract

Recognition of a pathogen avirulence (AVR) effector protein by a cognate plant resistance (R) protein triggers a set of immune responses that render the plant resistant. Pathogens can escape this so-called Effector-Triggered Immunity (ETI) by different mechanisms including the deletion or loss-of-function mutation of the *AVR* gene, the incorporation of point mutations that allow recognition to be evaded while maintaining virulence function, and the acquisition of new effectors that suppress AVR recognition. The Dothideomycete *Leptosphaeria maculans*, causal agent of oilseed rape stem canker, is one of the few fungal pathogens where suppression of ETI by an AVR effector has been demonstrated. Indeed, AvrLm4-7 suppresses Rlm3- and Rlm9- mediated resistance triggered by AvrLm3 and AvrLm5-9, respectively. The presence of *AvrLm4-7* does not impede *AvrLm3* and *AvrLm5-9* expression, and the three AVR proteins do not appear to physically interact. To decipher the epistatic interaction between these *L. maculans* AVR effectors, we determined the crystal structure of AvrLm5-9 and obtained a 3D model of AvrLm3, based on the crystal structure of Ecp11-1, a homologous AVR effector candidate from *Fulvia fulva*. Despite a lack of sequence similarity, AvrLm5-9 and AvrLm3 are structural analogues of AvrLm4-7 (structure previously characterized). Structure-informed sequence database searches identified a larger number of putative structural analogues among *L. maculans* effector candidates, including the AVR effector AvrLmS-Lep2, all produced during the early stages of oilseed rape infection, as well as among effector candidates from other phytopathogenic fungi. These structural analogues are named LARS (for Leptosphaeria AviRulence and Suppressing) effectors. Remarkably, transformants of *L. maculans* expressing one of these structural analogues, Ecp11-1, triggered oilseed rape immunity in several genotypes carrying *Rlm3*. Furthermore, this resistance could be suppressed by AvrLm4-7. These results suggest that Ecp11-1 shares a common activity with AvrLm3 within the host plant which is detected by Rlm3, or that the Ecp11-1 structure is sufficiently close to that of AvrLm3 to be recognized by Rlm3.

**Author summary:** An efficient strategy to control fungal diseases in the field is genetic control using resistant crop cultivars. Crop resistance mainly relies on gene-for-gene relationships between plant resistance (*R*) genes and pathogen avirulence (*AVR*) genes, as defined by Flor in the 1940s. However, such gene-for-gene relationships can increase in complexity over the course of plant-pathogen co-evolution. Resistance against the plant-pathogenic fungus *Leptosphaeria maculans* by *Brassica napus* and other *Brassica* species relies on the recognition of effector (AVR) proteins by R proteins; however, *L. maculans* produces an effector that suppresses a subset of these specific resistances. Using a protein structure approach, we revealed structural analogy between several of the resistance-triggering effectors, the resistance-suppressing effector, and effectors from other plant-pathogenic species in the Dothideomycetes and Sordariomycetes classes, defining a new family of effectors called LARS. Notably, cross-species expression of one LARS effector from *Fulvia fulva*, a pathogen of tomato, in *L. maculans* resulted in recognition by several resistant cultivars of oilseed rape. These results highlight the need to integrate knowledge on effector structures to improve resistance management and to develop broad-spectrum resistances for multi-pathogen control of diseases.

## Introduction

Fungi are the most devastating pathogens of plants, including crops of major economic importance. They represent a recurrent threat to agriculture and possess extreme adaptive abilities, resulting in the constant disease outbreaks [1,2]. Host invasion relies on effectors, key elements of pathogenesis, that modulate plant immunity and facilitate infection [3,4]. Fungal effector genes are diverse and typically encode small proteins, predicted to be secreted, with no or few homologues present in sequence databases, and an absence of known sequence motifs. In most phytopathogenic fungi, no large effector gene families have been identified [5]. Notably, effectors can have a dual role in plant-pathogen interactions, both targeting plant components and being targeted by plant resistance (R) proteins. Such dual-role effectors are known as avirulence (AVR) proteins because, in the presence of a corresponding R protein, they render the pathogen that produces them avirulent. Recognition of a pathogen AVR protein triggers a set of immune responses grouped under the term Effector-Triggered Immunity (ETI), frequently leading to a rapid localized cell death termed the hypersensitive response (HR) [6].

Breeding cultivars carrying *R* genes against pathogens is a common and powerful tool to control disease. However, the massive deployment of single *R* genes in the field exerts a strong selection pressure against avirulent pathogens that can become virulent through evolution of their *AVR* gene repertoire. Mechanisms leading to virulence include deletion, inactivation or down-regulation of the *AVR* gene, point mutations allowing recognition to be evaded while maintaining the virulence function of the AVR protein, or the acquisition of new effectors that suppress ETI [6–8]. Suppression of ETI by a fungal effector represents an efficient way to evade the selection pressure exerted by *R* genes in the field while maintaining the function of non-dispensable effectors. In some cases, the effector that suppresses ETI can itself be recognized by an R protein. A few examples of such strategies have been described in fungi [9–11], but the underlying mechanisms for the suppression of ETI by fungal effectors remain unexplained.

The Dothideomycete *Leptosphaeria maculans*, causal agent of oilseed rape stem canker or blackleg disease, is one of the fungal pathogens in which suppression of ETI by the presence of an *AVR* gene has been demonstrated. *L. maculans* can be controlled by combining qualitative and quantitative resistance of the host plant [12]. To date, ten *AVR* genes (called *AvrLm*) recognized by the products of *R* genes (called *Rlm*) from *Brassica napus* or other *Brassica* species have been identified [13–17] and share common characteristics: they encode small secreted proteins with no or low homologies in sequence databases, are located in repeat-rich regions of the genome, and are specifically expressed during the early stages of leaf infection. Among them, AvrLm4-7 suppresses Rlm3-mediated resistance triggered by AvrLm3 and Rlm9-mediated resistance triggered by AvrLm5-9 [9,14]. How AvrLm4-7 suppresses Rlm9-and Rlm3-mediated disease resistance is not known: the presence of *AvrLm4-7* does not impede *AvrLm3* and *AvrLm5-9* expression, and yeast two-hybrid (Y2H) assays suggest the absence of a physical interaction between AvrLm4-7, AvrLm5-9 and AvrLm3. While AvrLm5-9 and AvrLm3 share 29 % of amino acid sequence identity, a very low level of identity was found with AvrLm4-7 (15 %). AvrLm4-7 confers a dual recognition specificity by two distinct R proteins of oilseed rape, Rlm4 and Rlm7 [18], and loss of *AvrLm4-7* is associated with a fitness cost [19,20]. *AvrLm5-*9 and *AvrLm3*, on the other hand, are always present in *L. maculans* isolates, and only point mutation polymorphisms were reported, suggesting a high importance of these two effectors in pathogenicity towards *B. napus* [14,21,22]. Moreover, the silencing of *AvrLm3* led to a reduced aggressiveness [9].

Elucidation of the 3D structures of effectors may provide an effective strategy to resolve functional traits. Indeed, structure determination of effectors and the proteins with which they interact has provided key advances in our understanding of plant-pathogen interactions, including: the identification of protein functions that were not apparent from sequence analysis alone, the visualization of molecular interfaces of relevance to pathogen virulence and to plant immunity, and the identification of structural homologies in effectors that were not visible by sequence comparisons (reviewed in [23]). The crystal structure of AvrLm4-7 did not reveal similarities with documented effectors, but suggested a positively charged surface patch could be involved in AvrLm4-7 translocation into the cytoplasm of plant cells [24]. AvrLm4-7 escapes Rlm4-mediated recognition through a single point mutation [18] and Rlm7-mediated recognition through more drastic DNA changes (gene deletion, accumulation of mutations) or point mutations [25]. Blondeau et al. [24] identified a protein region involved in Rlm4-mediated recognition, and two regions involved in Rlm7-mediated recognition.

Here we describe the 3D structures of AvrLm5-9 and AvrLm3, whose recognition by Rlm9 and Rlm3, respectively, is masked by the presence of AvrLm4-7. Surprisingly, despite low sequence similarity, AvrLm5-9 and AvrLm3 are structural analogues of AvrLm4-7, sharing an anti-parallel β-sheet covered by α-helices. Structure-informed and pattern-based searches identified a larger number of putative structural analogues among AVR effectors and effector candidates of *L. maculans*, but also of other phytopathogenic fungi, including Ecp11-1 from the biotrophic tomato leaf mold fungus *Fulvia fulva* (formerly *Cladosporium fulvum*). Remarkably, transformants of *L. maculans* producing *F. fulva* Ecp11-1 triggered Rlm3*-* mediated immunity and this resistance could be suppressed by AvrLm4-7. These findings will enable hypotheses to be made about the way effectors suppress ETI and can guide recommendations on how to use plant R genes targeting AVRs belonging to structural families of effectors.

## Results

### Determination of the 3D structures of AvrLm5-9 and AvrLm3

To explore the putative molecular relationships between AvrLm4-7, AvrLm5-9 and AvrLm3, we set out to determine the 3D structures of the AvrLm3 and AvrLm5-9 effectors. AvrLm5-9 and AvrLm3 are rich in cysteines and therefore are difficult to express in soluble form in *Escherichia coli*. For the recombinant production of AvrLm5-9 and AvrLm3, we therefore chose the well-established *Pichia pastoris* eukaryotic expression system. The genes coding for the AvrLm5-9 and AvrLm3 proteins without their secretion signal peptides were cloned into expression vectors as fusion proteins with a purification His-tag and with thioredoxin. A TEV proteolytic cleavage site was inserted between thioredoxin and the effectors. The AvrLm5-9 fusion protein was well expressed and purified to homogeneity (Fig S1), but the yields of the AvrLm3 fusion protein were insufficient (*i.e*. about 50 mg of pure AvrLm5-9 per liter of cell culture against less than 1 mg of purified AvrLm3).

A small secreted protein with 37 % amino acid sequence identity with AvrLm3 was identified from *F. fulva* [26]. This protein, named Ecp11-1 (Extracellular protein 11-1), was found in apoplastic washing fluid samples harvested from compatible *F. fulva*–*Solanum lycopersicum* (tomato) interactions. Curiously, Ecp11-1 triggers an HR in multiple wild accessions of tomato. It is therefore likely that Ecp11-1 is an AVR effector recognized by a corresponding R protein (tentatively named Cf-Ecp11-1) in wild accessions of tomato [26]. We decided to produce Ecp11-1 using the same *P. pastoris*-based strategy as described for AvrLm5-9 and AvrLm3. The yields of Ecp11-1 production were sufficient to start structural studies (Fig S1, *i.e.* about 5 mg of purified Ecp11-1 per liter of cell culture).

After cleavage by the TEV protease and removal of thioredoxin, both AvrLm5-9 and Ecp11-1 provided good quality crystals. The structure of AvrLm5-9 was solved using iodine single wavelength anomalous diffusion (SAD) signal from a derivative crystal and then refined at 2.14 Å resolution using native data (Table S1). The crystallization liquor of AvrLm5-9 contained 80 mM of Ni^2+^ ions that proved mandatory for obtaining crystals. The structure revealed the presence of three Ni^2+^ ions that are involved in crystal contacts: two are found at the N- and C-termini of AvrLm5-9 as a ligand with two histidines from the native protein, by one histidine from a linker peptide and one from a crystal neighbor. The third Ni^2+^ ion is bound at the opposite end, and also interacts with histidines from two neighboring copies of AvrLm5-9 in the crystal. This suggests the bound Ni^2+^ ions are a crystallographic artefact. The complete AvrLm5-9 sequence could be placed into the electron density, which also accounted for two residues from the linker peptide at the N-terminus. The structure of AvrLm5-9 consists of a central β-sheet made of three anti-parallel β-strands (Fig 1A). An elongated peptide (residues 54 to 64) runs anti-parallel to β-strand 2, but only establishes a few main-chain H-bonds, and therefore is not categorized as a β-strand. One face of the β-sheet is covered by the long connections between the stands. The connection between the β1 and β2 strands is a curved α-helix and the connection between the β3 and β4 strands contains a shorter helix surrounded by two irregular peptide loops. AvrLm5-9 has three disulfide bridges, which are C^3^-C^119^, C^22^-C^69^ and C^26^-C^71^, named SS1, SS2 and SS3, respectively, according to protein numeration without the signal peptide (Fig 2) or C^22^-C^138^, C^41^-C^88^ and C^45^-C^90^ according to protein numeration with the signal peptide. The SS1 bridge knits the N- and C-termini together, while the other two disulfide bridges fix the long helix onto the β-sheet.

**Fig 1.**
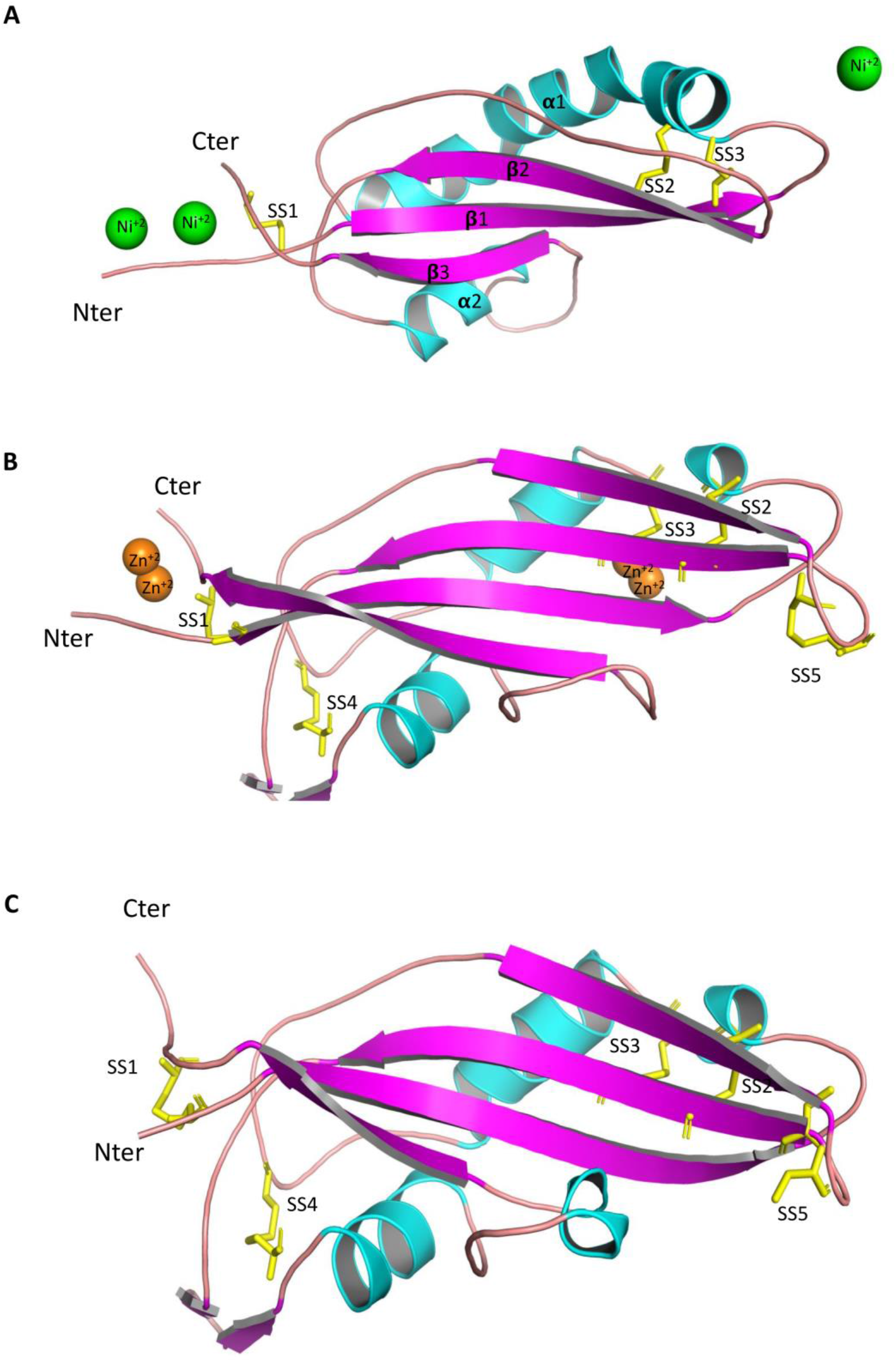
Crystal structures of AvrLm5-9 and Ecp11-1 and a 3D-model of AvrLm3. The proteins are represented as cartoons and coloured by secondary structure (α-helix in cyan, β-strand in magenta). N- and C-termini are labelled. The S-S bonds are represented by yellow sticks, and are numbered (numbering refers to sequence alignment in figure 2A). (A) Structure of AvrLm5-9. Ni^+2^ ions involved in crystal packing are shown as green spheres. (B) Structure of ECP11-1. Zn^+2^ ions involved in crystal packing are shown as orange spheres. (C) Swiss-Model structure of AvrLm3. The AvrLm3 amino acid sequence and the Ecp11-1 X-ray structure were used to compute the AvrLm3 three-dimensional model.

**Fig 2.**
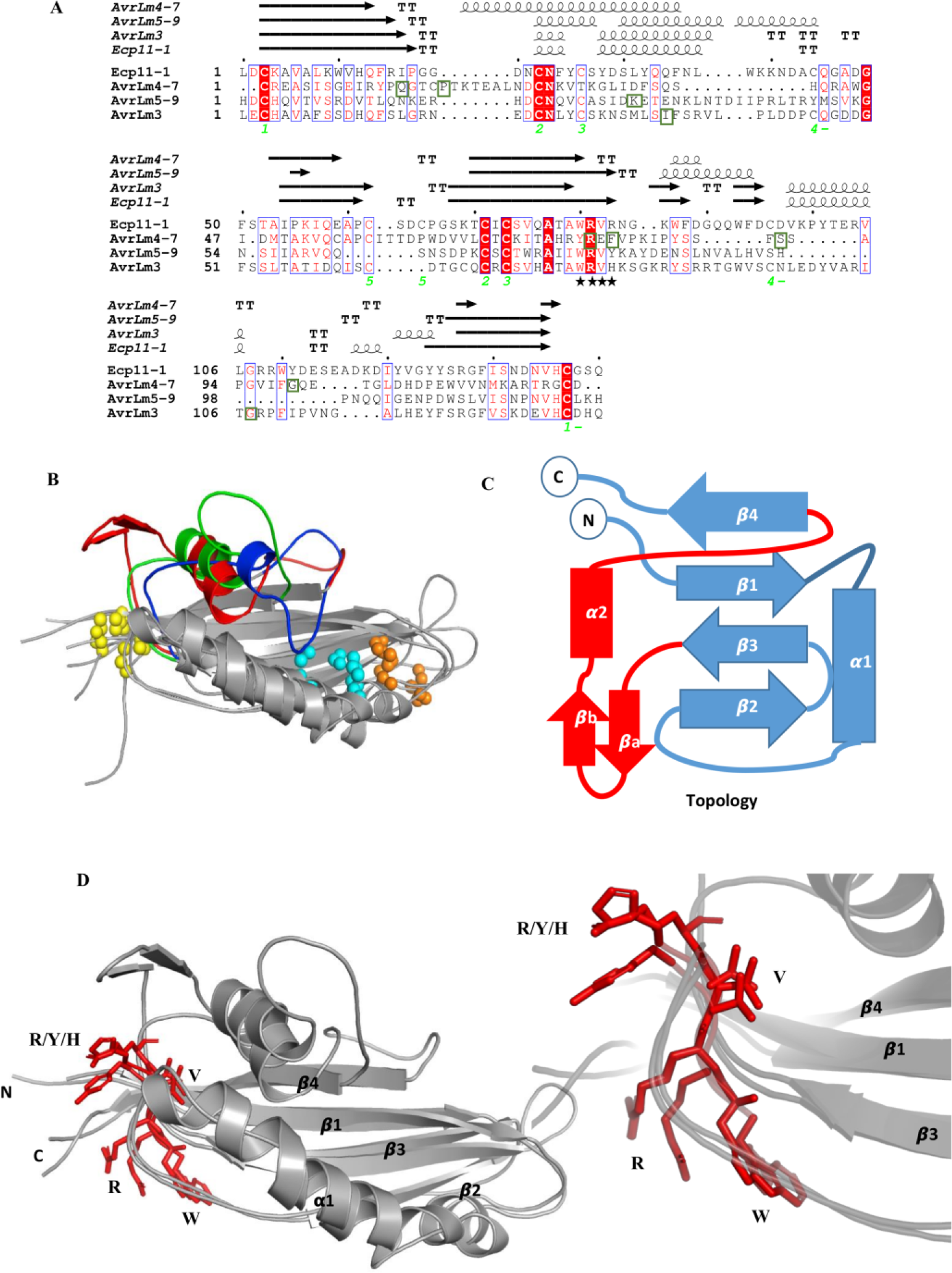
AvrLm4-7, AvrLm5-9, AvrLm3 and Ecp11-1 are structural analogues. (A) Structure-based protein sequence alignment of AvrLm4-7, AvrLm5-9, AvrLm3 and Ecp11-1. The S-S bridges of Ecp11-1 are labelled green and numbered to show their connectivity. Secondary structure for each protein (β-sheets, α-helices, and β-turns are rendered as arrows, squiggles and TT letters respectively) is shown above the alignment. Identical residues are in red boxes and similar residues are in red. The conserved motif WR(F/L/V)(R/K) is labeled by black stars. The residues for whom mutations are associated with a switch to virulence are in green boxes. The Figure was made with the ESPript server [27]. (B) Superposition of AvrLm4-7, AvrLm5-9 and AvrLm3. The variable connections between β3 and β4 are coloured in red (AvrLm3), blue (AvrLm4-7) and green (AvrLm5-9). The conserved or neighbouring S-S bridges are indicated as yellow, orange and cyan spheres (indicated as 1, 2 and 3, respectively, in panel A). (C) Representation of conserved topology, with the variable connexion between β3 and β4 coloured in red. (D) Superposition of AvrLm5-9, Ecp11-1 and AvrLm3 (all in grey), with their conserved WR(F/L/V)(R/K) motif at the exit of β3 represented by red sticks. A zoom on the motif is represented in the lower right corner.

The presence of a metal, in this case Zn^2+^, was also mandatory for the crystallization of Ecp11-1. Speculating that the Ecp11-1 crystal would also bind a bivalent ion, we successfully solved its structure at 1.6 Å resolution by using the SAD-signal of Zn^2+^. The crystals contained one copy of Ecp11-1 in the asymmetric unit. The complete sequence could be positioned into the electron density. The analysis of the anomalous signal revealed the presence of two Zn-clusters in the structure that resemble those of the Ni-ions in AvrLm5-9. One Zn-cluster is bound near the N- and C-termini and involves histidines from the native protein and from the linker peptide and side chains from a crystal neighbor. Another Zn-cluster is found at the opposite end and is composed by residues emanating from neighboring Ecp11-1 molecules. Ecp11-1 forms an anti-parallel four-stranded β-sheet with (+2,-1,-2) topology (Fig 1B and 2C). Strands β1 and β2 are connected by a peptide composed of a helical turn and a short helix surrounded by irregular peptides. The long connection between strands β3 and β4 contains a β-hairpin, a short helix and some irregular peptide stretches. Ecp11-1 contains five disulfide bridges: C^3^-C^137^ (named SS1) that connects the N- and C-termini, C^22^-C^71^ (named SS2) and C^26^-C^73^ (named SS3) that link the β1-β2 connection to strand β3. The two remaining disulfide bridges (C^44^-C^96^ and C^62^-C^65^, named SS4 and SS5) (according to protein numbering in Fig 2) are found in the irregular loop regions.

AvrLm3 and Ecp11-1 share 37 % identity and 59% similarity, and there are only minor insertions/deletions between the two sequences. We consequently constructed a 3D-model of AvrLm3 using the Swiss-Model web server (https://swissmodel.expasy.org/). The sequence alignment and structure superposition show conservation of all 10 cysteines, suggesting both proteins form an identical set of disulfide bridges (Fig 1B and C).

### AvrLm4-7, AvrLm5-9, AvrLm3 and Ecp11-1 form a structural family

The sequence similarity between AvrLm5-9 and Ecp11-1/AvrLm3 is weak. The sequence similarity between AvrLm4-7 on the one hand and AvrLm5-9 and Ecp11-1 on the other is even weaker than between the latter two (Fig.2A-B). Nevertheless the 3D structures of all these effectors are strongly related: AvrLm5-9 and Ecp11-1/AvrLm3 share the same fold with AvrLm4-7 (Fig 2B), while it displays suppressive interactions with AvrLm5-9 and AvrLm3. Both AvrLm4-7 (β4) and AvrLm5-9 (β2) have an irregular β-strand, but these are oriented and positioned in the same way as the corresponding strands in Ecp11-1. To fully appreciate the similarities between the three effectors, we superposed their structures using the DALI webserver (http://ekhidna2.biocenter.helsinki.fi/dali/). Superposition of Ecp11-1 and AvrLm5-9 gives a Z-score of 8.9 and an RMSD value of 3.3 Å (106 residues aligned) and superposition of Ecp11-1 and AvrLm4-7 gives a Z-score of 5.6 and an RMSD value of 3.8 Å (106 residues aligned). The main differences between the three proteins are found in the regions connecting the strands (Fig 2B and 2C). The structures of the three effectors are stabilized by disulfide bridges, two of which are overlapping in the three proteins. All three effectors have a disulfide bridge that connects the N- and C-termini, and also at least one disulfide bridge that links the helical connection between β1-β2 and strand β3.

A search in the Protein Data Bank (PDB) for structural analogues of Ecp11-1 and AvrLm5-9 using the DALI webserver returned AvrLm4-7 and the yeast elongation factor 1B. The latter protein (100 residues) indeed has the same β-sheet topology and similar connections, but shares no significant sequence identity with the effectors, has no disulfide bridges and the region of EF1B relevant for its function is not conserved in Ecp11-1 and AvrLm5-9. The similarity is therefore probably not biologically relevant.

### Polymorphic residues identified in *L. maculans* and *F. fulva* populations are mainly located on loop regions of AvrLm5-9, AvrLm3 and Ecp11-1

We then set out to identify regions putatively required for recognition by cognate R proteins by mapping polymorphic residues onto their respective structures. We therefore exploited previous population studies that had reported polymorphisms in AvrLm5-9, AvrLm3 and Ecp11-1. Only three polymorphic residues of AvrLm5-9 were reported in *L. maculans* populations [14,21]. One event caused a switch to virulence towards both *Rlm5* and *Rlm9* (R^29^ to stop), a point mutation at residue R^55^ to T or K leads to virulence towards Rlm9 (K^36^ according to the protein version and numbering in Fig 2, since the AvrLm5-9 protein whose structure was determined conferred virulence towards Rlm9). A third polymorphism had no effect on the interaction with Rlm5 or Rlm9 (R^38^ to L) (R^19^ according to protein numbering in Fig 2). The stop mutation evidently leads to a truncated and non-functional protein. The two last polymorphic residues were mapped onto the AvrLm5-9 3D structure (Fig 3A). The mutation leading to virulence towards *Rlm9* is situated in the middle of the long helix, and the mutation having no effect on the interaction with Rlm9 and Rlm5 is in a loop connecting β1 and β2. Neither mutation is expected to perturb the 3D structure.

**Fig 3.**
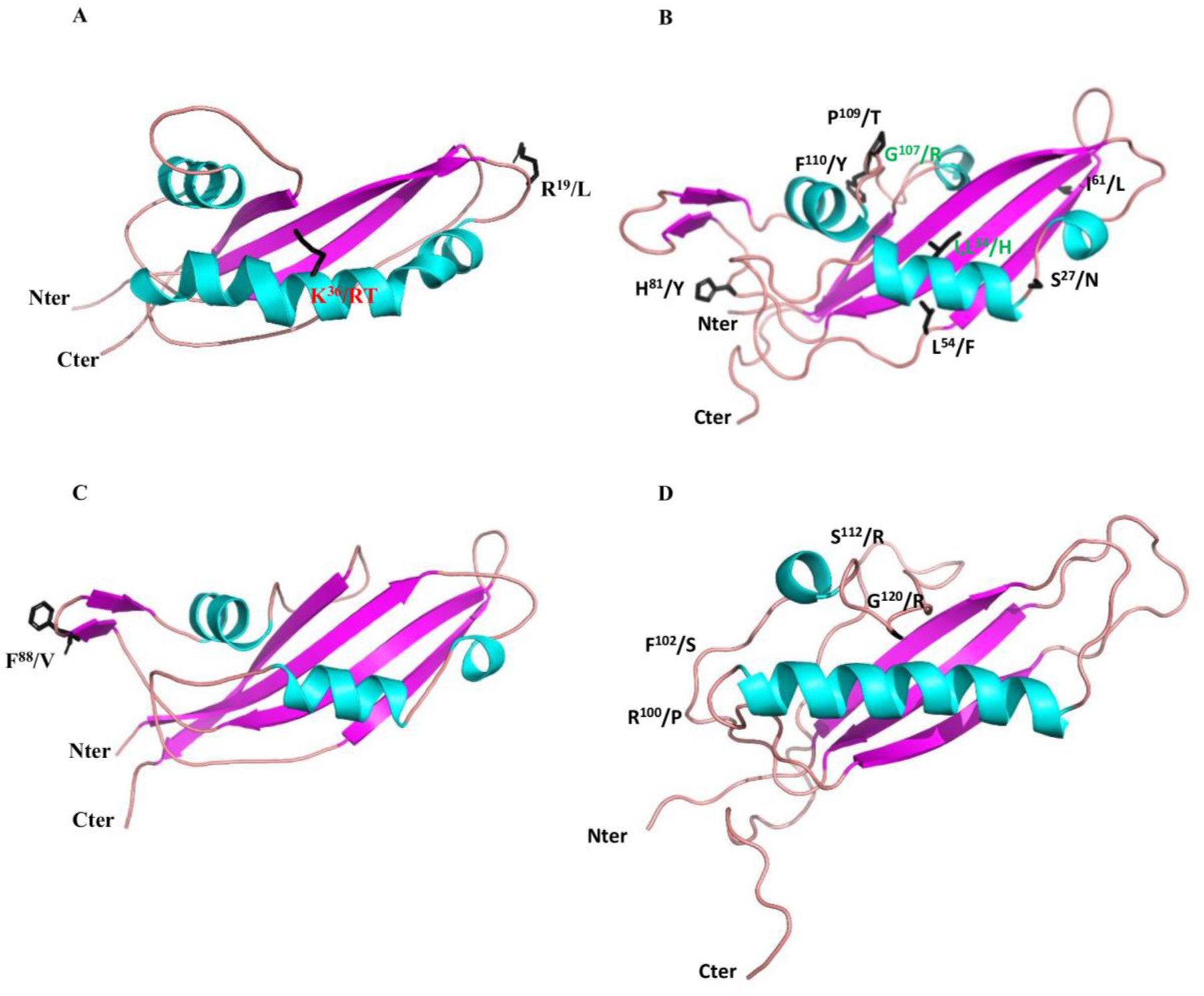
Localisation of polymorphic residues on the 3D structures of AvrLm5-9, AvrLm3, Ecp11-1 and AvrLm4-7. Polymorphic residues were identified in populations of *L. maculans* ‘brassicae’ and *F. fulva.* (A) Positions of polymorphisms in AvrLm5-9 [14,21]. Amino acid change involved in virulence toward *Rlm9* is indicated in red. (B) Positions of polymorphisms in AvrLm3 [22]. Amino acid changes shared among isolates virulent toward *Rlm3* are indicated in green. (C) Polymorphism in Ecp11-1 [26]. It is currently unclear whether this variant impacts protein function. (D) Polymorphic positions in AvrLm4-7 leading to virulence towards Rlm4 or Rlm7 [18,24,25].

Sequencing of AvrLm3 in a collection of worldwide *L. maculans* isolates defined 22 different alleles, corresponding to 14 non-synonymous mutations leading to 11 isoforms of the protein [22]. We projected eight of the 11 polymorphic amino acids onto the AvrLm3 3D structure (the remaining three are located in the signal peptide and thus could not be plotted) (Fig 3B). Most of the AvrLm3 variants concern amino acid positions with surface-exposed side chains, and none of the variants affects the cysteines. Only one polymorphic substitution (L^78^ to F) (L^54^ according to protein numbering in Fig 2) is found in a residue involved in hydrophobic packing, but the mutation is conservative. These observations suggest that all alleles should lead to well-folded proteins. Six of the 11 AvrLm3 isoforms were only found in isolates avirulent towards *Rlm3*. These amino acids are scattered over the surface of the protein. Only two amino acid residues consistently differed between virulent and avirulent isoforms: I/L^58^H and G^131^R (I^34^ and G^107^ according to protein numbering in Fig 2). Of interest, amino acid residue I^58^ is located in the same region of the structure in AvrLm3 as residue K^55^ (responsible for the switch to virulence towards Rlm9) in the AvrLm5-9 structure. Similarly, residue G^131^ in AvrLm3 is located in the same region as residue G/R^120^ in AvrLm4-7 (responsible for the switch to virulence towards Rlm4 [18]; Fig 3D; G^99^according to Fig 2).

Interestingly, substitutions at I^34^ and G^107^ are always coupled and likely are responsible for the virulent phenotype towards *Rlm3*. They are positioned in the regions connecting the strands, and the folding brings them relatively close together in space, suggesting this site could be important for its function. Remarkably, the positions in the 3D structures of the polymorphic residues G^131^R (AvrLm3, G^107^ in Fig 3) and G^120^R (AvrLm4-7; [24]) are very similar, suggesting the same protein regions could be involved in the virulence phenotype.

From a population study on a worldwide collection of *F. fulva* strains [26], a single non-synonymous F^119^V substitution was identified in Ecp11-1. W^118^, F^119^ and W^124^ (W^87^, F^88^ and W^93^ according to protein numbering in Fig 2) form an exposed hydrophobic surface patch contiguous to a conserved WR(F/L/V)(R/K) motif (see below for more information on this motif). It is not yet known whether this non-synonymous substitution affects the ability of Ecp11-1 to trigger an HR in tomato plants carrying the putative Cf-Ecp11-1 R protein (Fig 3C).

### Structural analogues of AvrLm4-7, AvrLm5-9 and Ecp11-1 are present in Leptosphaeria maculans

The strong structural similarities between AvrLm4-7, AvrLm5-9, AvrLm3 and Ecp11-1 suggest that these four proteins belong to a fungal effector family, characterized by a four-stranded β-sheet, an α-helix and three conserved disulfide bridges (Fig 2A and 2C). To identify other structurally related family members, a hidden Markov model (HMM)-based profile search was performed on the predicted protein repertoire from *L. maculans* (v23.1.3 isolate). An iterative search was carried out with each effector structure, using a cut-off E-value of 1 and an overlap cut-off of 50%. A total of three iterations were performed. At each iteration, proteins longer than 160 amino acids were removed. Seven potential structural analogues were found for AvrLm4-7, six for Ecp11-1 and four for AvrLm5-9. The search with Ecp11-1 retrieved both AvrLm3 and AvrLm4-7, the remaining four were also found with the search using AvrLm4-7. The search with AvrLm5-9 retrieved AvrLm3 together with other candidates that were not found with AvrLm4-7 or Ecp11-1, including the AVR protein AvrLmS-Lep2 (Table 1; [16]). A total of thirteen structural analogues were thus identified in *L. maculans* (Table 1).

**Table 1.**
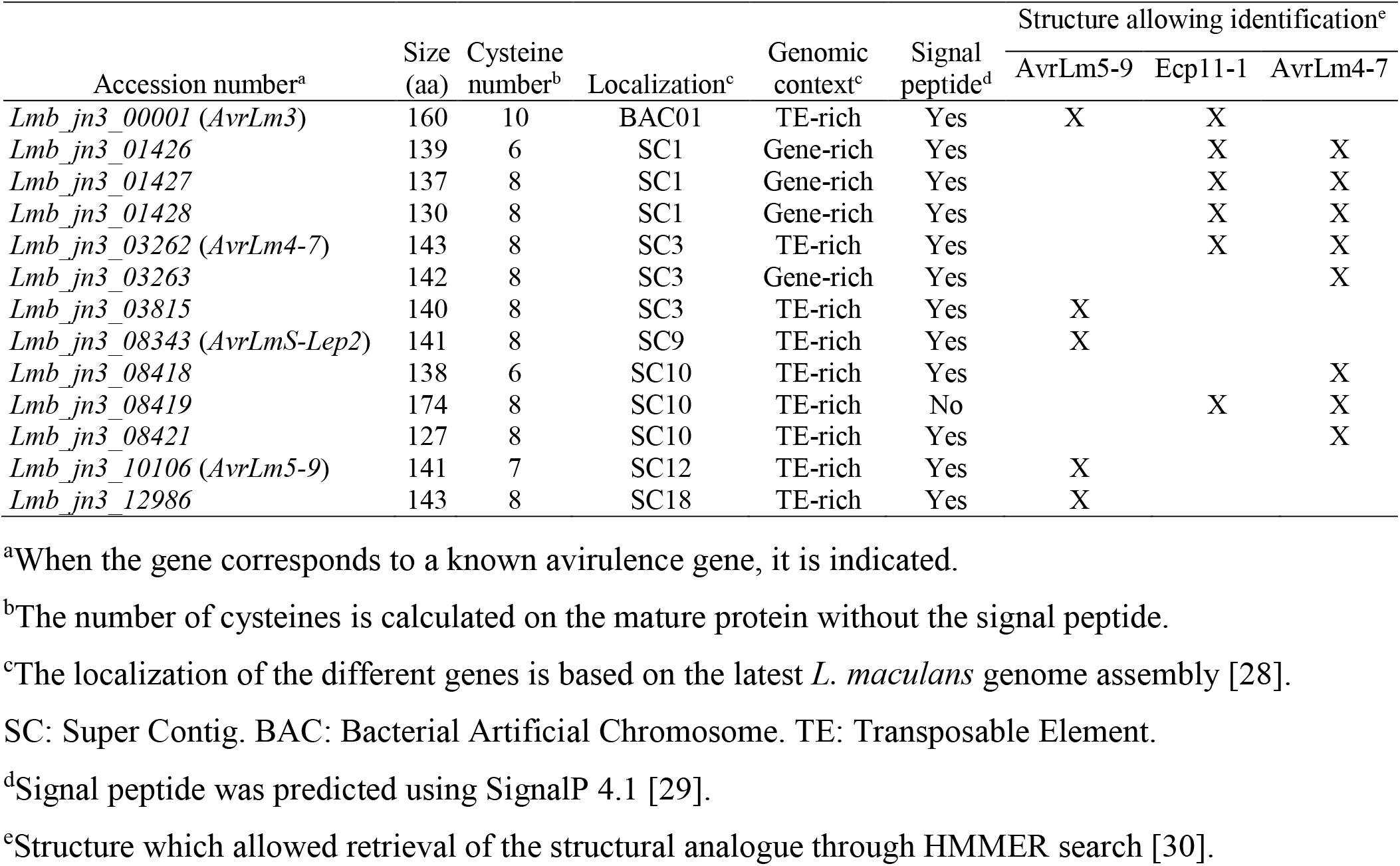
Characteristics of the structural analogues identified in *L. maculans*.

| Accession number<sup>a</sup> | Size<br>(aa) | Cysteine<br>number<sup>b</sup> | Localization<sup>c</sup> | Genomic<br>context<sup>c</sup> | Signal<br>peptide<sup>d</sup> | Structure allowing identification<sup>e</sup> |  |  |
| --- | --- | --- | --- | --- | --- | --- | --- | --- |
|  |  |  |  |  |  | AvrLm5-9 | Ecp11-1 | AvrLm4-7 |
| *Lmb_jn3_00001* (*AvrLm3*) | 160 | 10 | BAC01 | TE-rich | Yes | X | X |  |
| *Lmb_jn3_01426* | 139 | 6 | SC1 | Gene-rich | Yes |  | X | X |
| *Lmb_jn3_01427* | 137 | 8 | SC1 | Gene-rich | Yes |  | X | X |
| *Lmb_jn3_01428* | 130 | 8 | SC1 | Gene-rich | Yes |  | X | X |
| *Lmb_jn3_03262* (*AvrLm4-7*) | 143 | 8 | SC3 | TE-rich | Yes |  | X | X |
| *Lmb_jn3_03263* | 142 | 8 | SC3 | Gene-rich | Yes |  |  | X |
| *Lmb_jn3_03815* | 140 | 8 | SC3 | TE-rich | Yes | X |  |  |
| *Lmb_jn3_08343* (*AvrLmS-Lep2*) | 141 | 8 | SC9 | TE-rich | Yes | X |  |  |
| *Lmb_jn3_08418* | 138 | 6 | SC10 | TE-rich | Yes |  |  | X |
| *Lmb_jn3_08419* | 174 | 8 | SC10 | TE-rich | No |  | X | X |
| *Lmb_jn3_08421* | 127 | 8 | SC10 | TE-rich | Yes |  |  | X |
| *Lmb_jn3_10106* (*AvrLm5-9*) | 141 | 7 | SC12 | TE-rich | Yes | X |  |  |
| *Lmb_jn3_12986* | 143 | 8 | SC18 | TE-rich | Yes | X |  |  |

<sup>a</sup>When the gene corresponds to a known avirulence gene, it is indicated.
<sup>b</sup>The number of cysteines is calculated on the mature protein without the signal peptide.
<sup>c</sup>The localization of the different genes is based on the latest *L. maculans* genome assembly [28].
SC: Super Contig. BAC: Bacterial Artificial Chromosome. TE: Transposable Element.
<sup>d</sup>Signal peptide was predicted using SignalP 4.1 [29].
<sup>e</sup>Structure which allowed retrieval of the structural analogue through HMMER search [30].

We then wanted to confirm that the retrieved protein analogues have similar structures. We therefore constructed 3D structural models of the different analogues found by the HMM search using the alphaFold program via the colab server (https://colab.research.google.com/github/sokrypton/ColabFold/blob/main/AlphaFold2.ipynb; [31]). For all retrieved sequence analogues, alphaFold proposed models that had the same fold as AvrLm4-7 (with the exception of Lmb_jn3_08343 and Lmb_jn3_12986 for which no reliable models could be proposed; Fig S3). Moreover, the cysteine bridges in all these models were superposable. We conclude that these sequences possess the same fold characterized by an antiparallel β-sheet and a characteristic set of disulfide bonds. This suggests they form a homologous family that we will name from this point forward as the LARS (for Leptosphaeria AviRulence and Suppressing) effector family.

### LARS effectors of *L. maculans* share common characteristics and are expressed *in planta*

The characteristics of LARS structural analogues identified in *L. maculans* using a HMM search are summarized in Table 1. The genomic location of the genes encoding structural analogues was investigated using the latest assembly of the *L. maculans* genome [28]. The majority of these genes are located in genomic regions rich in remnants of transposable elements, with the exception of a group of three neighboring genes (*Lmb_jn3_01426*, *Lmb_jn3_01427* and *Lmb_jn3_01428*), located in a gene-rich region, and a gene that had been previously described as a paralogue of *AvrLm4-7* (*Lmb_jn3_03263*; [18]), which is located at the border of a gene-rich region. These genes are mostly located on different Super Contigs. However, several genes are neighbours. This is the case for (i) *Lmb_jn3_01426*, *Lmb_jn3_01427* and *Lmb_jn3_01428*; (ii) *Lmb_jn3_08418*, *Lmb_jn3_08419* and *Lmb_jn3_08421*; (iii) *Lmb_jn3_03262* (*AvrLm4-7*) and *Lmb_jn3_03263*. The average size of these proteins is 140 amino acids, and they all have between six and 10 cysteines. For the whole family, apart from Lmb_jn3_08419, we were able to predict a signal peptide, suggesting that these proteins are secreted by the fungus.

RNAseq data corresponding to different stages of infection of oilseed rape cotyledons by *L. maculans* as well as during axenic growth of *L. maculans* on V8 solid medium were recently generated [32], and expression kinetics of the LARS effector genes analyzed (Fig 4). All the genes showed over-expression during the biotrophic / asymptomatic phase of cotyledon infection. A closer examination of the expression profiles distinguished three different patterns.

**Fig 4.**
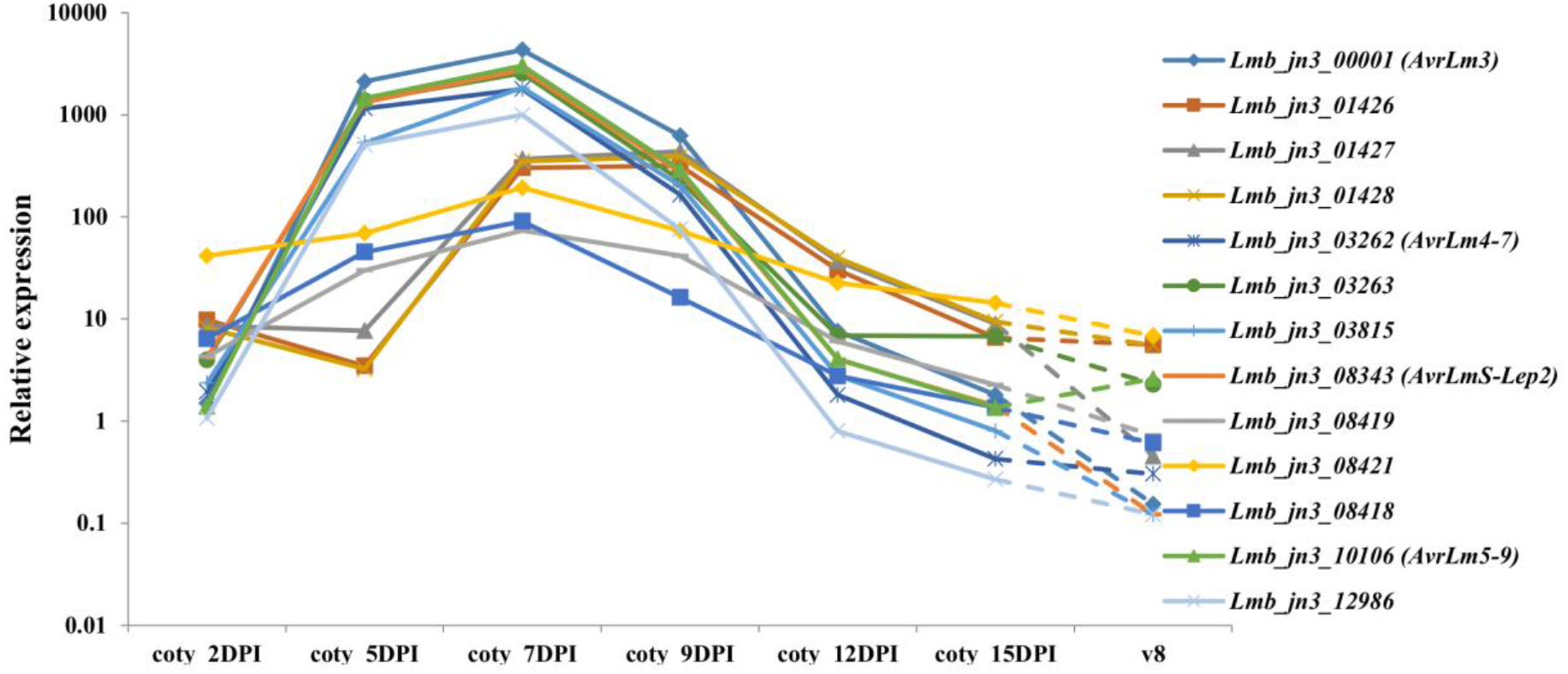
Expression kinetics of the LARS structural analogues identified in *L. maculans* ‘brassicae’. Expression is based on RNAseq data described in [32] normalized by the total number of sequences per condition, counts per million (CPM), and represented on a logarithmic scale, following inoculation of cotyledons from a susceptible oilseed rape cultivar, Darmor-Bzh. Coty (cotyledons of the susceptible oilseed rape cultivar, Darmor-Bzh infected by *L. maculans ‘*brassicae*’* 2, 5, 7, 9, 12 and 15 days post-inoculation (DPI)); v8 (growth on solid medium V8 for one week).

The first pattern represented genes that were highly expressed as early as 5 days post-inoculation(DPI) with a peak at 7 DPI, and included the *AVR* genes of the family: *Lmb_jn3_00001* (*AvrLm3*), *Lmb_jn3_03262* (*AvrLm4-7*), *Lmb_jn3_03263*, *Lmb_jn3_03815*, *Lmb_jn3_08343* (*AvrLmS-Lep2*), *Lmb_jn3_10106* (*AvrLm5-9*) and *Lmb_jn3_12986*. The second pattern of genes was characterized by a low expression at 5 DPI and a plateau between 7 and 9 DPI: *Lmb_jn3_01426*, *Lmb_jn3_01427* and *Lmb_jn3_01428*. The last pattern grouped genes whose expression peaked lower at 7 days post-inoculation: *Lmb_jn3_08418*, *Lmb_jn3_08419* and *Lmb_jn3_08421*. Of interest, genes with the same expression profile are neighbors in the *L. maculans* genome. Finally, on V8 medium, all the genes were very lowly expressed, which indicates that they are overexpressed during the primary infection of oilseed rape by *L. maculans*.

### LARS effectors are present in other phytopathogenic fungi

To find out whether the LARS effector family has members in other fungi, a new HMM-based profile search was performed on an in-house database, composed of annotated proteomes of 163 fungal strains, corresponding to 116 species, mostly Dothideomycetes and Sordariomycetes with contrasting lifestyles (phytopathogens, entomopathogens, endophytes, saprophytes, mycoparasites, Table S3). The in-house database was iteratively searched with each effector structure file, using a cut-off E-value of 1 and a cut-off overlap of 50 %. At each iteration, proteins longer than 160 amino acids were removed. This HMM-search identified, after three iterations, 34 potential structural analogues using AvrLm4-7, 32 with Ecp11-1 and two using AvrLm5-9. Interestingly, Ecp11-1 was found using AvrLm5-9 as a template, but not using AvrLm4-7. Combined with the analogues found in *L. maculans* ‘brassicae’, 49 non-redundant proteins were identified (Fig 5). These potential structural analogues originate from 13 fungal species, with the majority from species closely related to *L. maculans* ‘brassicae’ (*L. maculans ‘*lepidii’, *L. biglobosa* ‘brassicae’ and *L. biglobosa ‘*thlaspii’), Dothideomycetes (*Macrophomina phaseolina*, *Pyrenophora tritici-repentis*, *P. teres*, *Corynespora cassiicola*, *Stemphylium lycopersici* and *F. fulva*), and a few Sordariomycetes (*Colletotrichum orbiculare, C. gloeosporioides* and *C. higginsianum*; Fig 5A, Fig S2 and Table S3).

**Fig 5.**
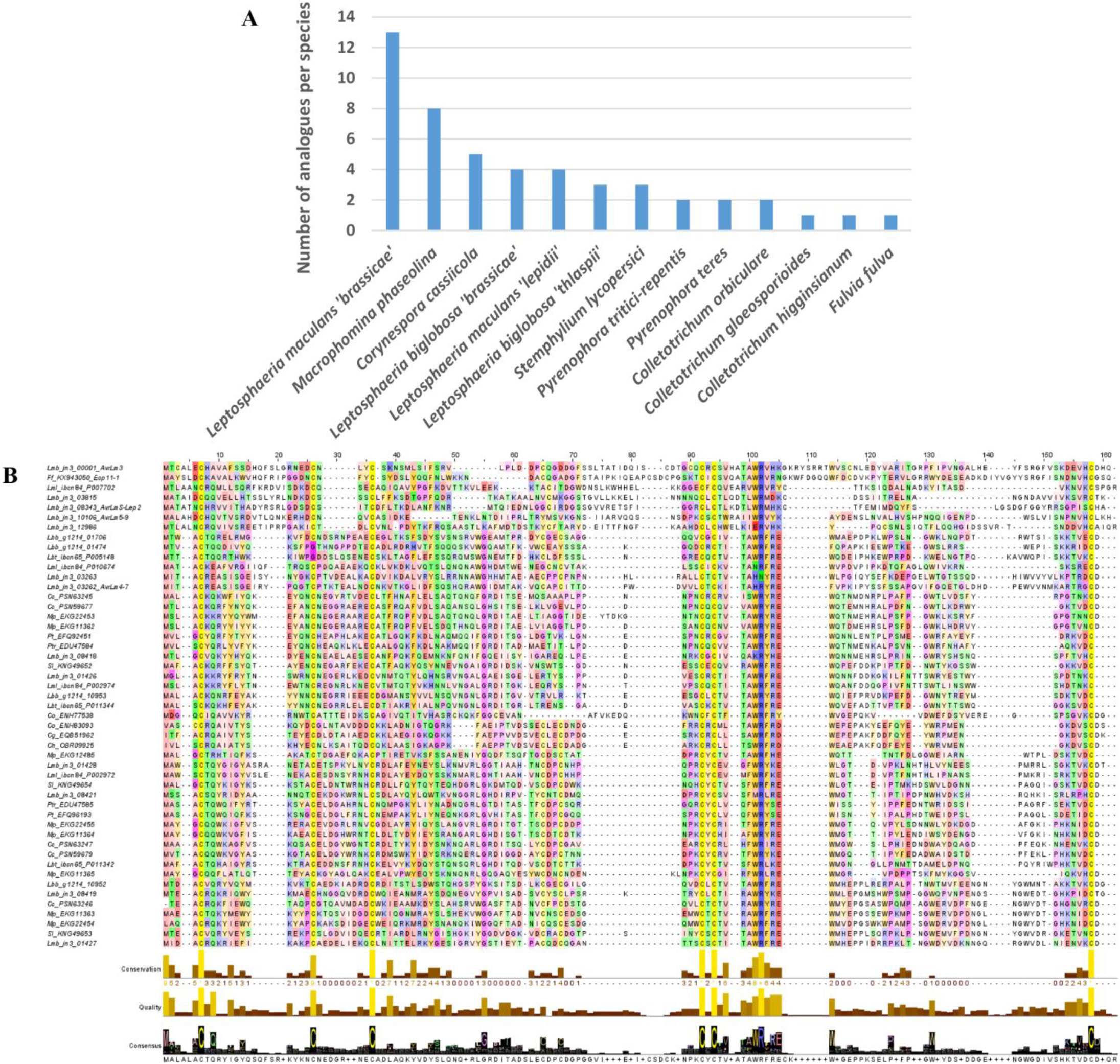
Identification of LARS effector structural analogues in *L. maculans* ‘brassicae’ and other phytopathogenic fungi. (A) Species distribution of number of structural analogues of AvrLm4-7, AvrLm5-9 and Ecp11-1 found with a low-stringency HMM search on a database encompassing 163 predicted proteomes of 116 fungal species. (B) Multiple sequence alignment of the 50 potential structural analogues and the sequences of the known structures (AvrLm4-7=4fprA / AvrLm5-9=0a59A / Ecp11-1=0cp1A). The secondary structures of the latter were calculated using the software STRIDE and were added at the bottom of the alignment (H=helix, G=3-10 helix, E= β strand, B=β bridge, C=coil, T=turn) above the residue conservation measure, the local alignment quality and the consensus logo. The alignment was displayed using the software Jalview. In the displayed alignment, amino acids which do not align with any of the three sequences AvrLm4-7, AvrLm5-9 and Ecp11-1 have been removed.

A sequence logo derived from the multiple alignment of the structural analogues highlighted conserved features of the LARS effectors (Fig 5B). Although the number of cysteines is variable between the different members, six cysteines can be aligned between the majority of the LARS members. The cysteines near the N- and C-termini form a disulfide bridge in all available structures, and this is likely the case for all of the proteins identified. The remaining aligned cysteines are not always involved in superimposable disulfide bridges in the three available structures. All structures have a putative disulfide bridge that connects α-helix 1 to β-strand 3. The third conserved cysteine pair establishes a different disulfide bridge, however, in AvrLm4-7 on the one hand and AvrLm5-9, Ecp11-1 and AvrLm3 on the other (Fig 2A). Apart from the cysteines, a WR(F/L/V)(R/K) sequence motif is very well conserved in all sequences (Fig 2D), positioned at the exit of the third β strand. The motif on one side crosses the disulfide bridge that connects the N- and C-termini, and on the other lies against the α–loop that precedes the second strand. Residues of this strand are less well conserved.

Cg, Colletotrichum gloeosporioides; Ch, Colletotrichum higginsianum; Co, Colletotrichum orbiculare; Cc, Corynespora cassiicola; Ff, Fulvia fulva; Lbb, Leptosphaeria biglobosa ‘brassicae’; Lbt, Leptosphaeria biglobosa ‘thlaspii’; Lmb, Leptosphaeria maculans ‘brassicae’; Lml, Leptosphaeria maculans ‘lepidii’; Mp, Macrophomina phaseolina; Pt, Pyrenophora teres; Ptr, Pyrenophora tritici-repentis; Sl, Stemphylium lycopersici Remarkably, the tryptophan residue of the motif is fully exposed to the solvent. The F/L/V residue fits into the hydrophobic core of the protein. The conservation of the motif and its exposure on the surface suggest that it might be involved in a functional interaction.

### Mutations in the WR(F/L/V)(R/K) conserved motif in AvrLm4-7 contribute to abolish its ability to suppress Rlm3-mediated recognition, but only when AvrLm4-7 escapes both Rlm7 and Rlm4-mediated recognition

We investigated whether the WR(F/L/V)(R/K) sequence motif, which is well conserved within the LARS structural family, could be involved in the ability of AvrLm4-7 to suppress recognition of AvrLm3 by Rlm3. Previous studies performed site-directed mutagenesis on AvrLm4-7 residues to investigate its ability to suppress the recognition of AvrLm3 by Rlm3 and to trigger Rlm7 and Rlm4-mediated immunity [22,24]. We extended these data with additional mutagenesis experiments (Table 2 and Fig S4). Mutagenesis was performed on an allele of AvrLm4-7 conferring both Rlm7 and Rlm4-mediated recognition or only Rlm7-mediated recognition (G^120^R mutation). Mutations R^100^P and F^102^S led to a switch to virulence towards *Rlm7* cultivar and abolished the ability of AvrLm4-7 to suppress Rlm3-mediated recognition of AvrLm3, but only in a G^120^R context, suggesting that both R^100^ or F^102^ and G^120^ are necessary to mask AvrLm3 recognition and induce *Rlm7* immunity. In contrast, mutation S^112^R, located close to G^120^, was sufficient to escape Rlm4 and Rlm7-mediated recognition and abolish the ability of AvrLm4-7 to suppress Rlm3-mediated recognition of AvrLm3.

**Table 2:** Interaction phenotypes on *Brassica napus* resistant genotypes of *Leptosphaeria maculans* wild-type or transformed isolates with different site-directed mutagenized alleles of *AvrLm4-7*.

| Isolate | Interaction Phenotype on |  |  | Reference |
| --- | --- | --- | --- | --- |
|  | *Rlm3* | *Rlm4* | *Rlm7* |  |
| G06-E107 | A | V | V | [25] |
| G06-E107-AvrLm4-7 | V | A | A | [22] |
| G06-E107-AvrLm4-7 G<sup>120</sup>R | V | V | A | [22] |
| G06-E107-AvrLm4-7R<sup>100</sup>P | V | A | A | This study |
| G06-E107-AvrLm4-7R<sup>100</sup>P G<sup>120</sup>R | A | V | V | [22,24] |
| G06-E107-AvrLm4-7 F<sup>102</sup>S | V | A | A | This study |
| G06-E107-AvrLm4-7 F<sup>102</sup>S G<sup>120</sup>R | A | V | V | [24]; this study |
| G06-E107-AvrLm4-7S<sup>112</sup>R | A | V | V | This study |
| G06-E107-AvrLm4-7S<sup>112</sup>R G<sup>120</sup>R | A | V | V | [22,24] |

### A structural analogue from *F. fulva* triggers recognition by oilseed rape R protein Rlm3, and this recognition is masked by the presence of AvrLm4-7

In an attempt to determine whether members of the LARS family from other phytopathogenic fungi are recognized by R proteins from oilseed rape and whether that recognition could be masked by AvrLm4-7, we set out to determine whether Rlm3 can recognize Ecp11-1 of *F. fulva*, and if so, whether AvrLm4-7 can ‘hide’ Ecp11-1 from Rlm3-mediated recognition.

As a first stage, a construct containing the *ECP11-1* coding sequence and terminator under the control of the *AvrLm4-7* promoter was introduced via *Agrobacterium tumefaciens*-mediated transformation into two isolates virulent towards *Rlm3*, *Rlm4* and *Rlm7*: Nz-T4 and v45.15. The corresponding transformants (10 Nz-T4-*ECP11-1* and 9 v45.15-*ECP11-1*) were inoculated onto *B. napus* cvs Pixel (*Rlm4*) and 15.22.4.1 (*Rlm3*; Fig 6). All transformants, as the wild type isolates, were virulent on Pixel. In contrast, and differing from their parental isolates, all transformants were avirulent on 15.22.4.1. We conclude that Ecp11-1, like AvrLm3, can be recognized by Rlm3. To confirm this result, we inoculated five transformants on two additional and unrelated oilseed rape cultivars carrying *Rlm3* (Grizzly and Columbus) and confirmed a resistance phenotype on these cultivars triggered by Ecp11-1 (Fig S5). Moreover, we also inoculated the transformants on the oilseed rape line 15.23.4.1, a sister line of 15.22.4.1 also issued from individual plants from cv. Rangi, carrying *Rlm7* instead of *Rlm3*, and obtained a susceptibility phenotype (Fig S5). These results support our conclusions that Ecp11-1 can be recognized by Rlm3.

**Fig 6.**
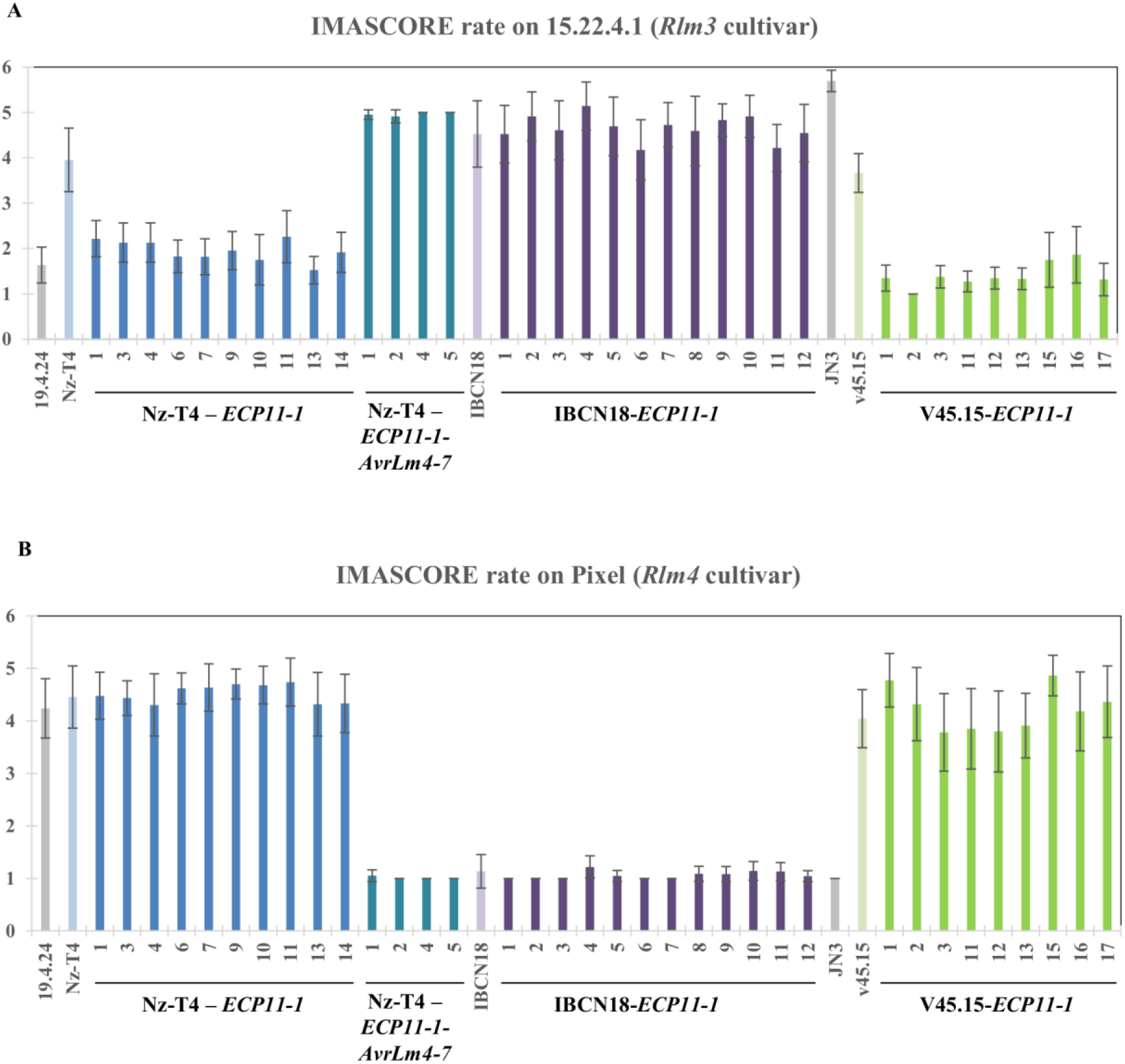
Ecp11-1 of *F. fulva* triggers Rlm3-mediated recognition in *L. maculans*, masked by AvrLm4-7. Wild type isolates Nz-T4 (a3a4a7), IBCN18 (a3A4A7) and v45.15 (a3a4a7), as well as Nz-T4, IBCN18 and v45.15 transformants carrying *ECP11-1*, and Nz-T4 transformants carrying both *ECP11-1* and *AvrLm4-7* were inoculated onto cotyledons of a cultivar carrying *Rlm3* (15.22.4.1, A) or *Rlm4* (Pixel, B). Pathogenicity was measured 13 days post-inoculation. Results are expressed as a mean scoring using the IMASCORE rating comprising six infection classes (IC), where IC1 to IC3 correspond to resistance, and IC4 to IC6 to susceptibility [33]. Error bars indicate the standard deviation of technical replicates. 19.4.24 (A3a4a7) and JN3 (a3A4A7) were used as controls of the AvrLm3/Rlm3, and AvrLm4-7/Rlm4 and Rlm7 interaction phenotypes, respectively.

At a second stage, we introduced *ECP11-1* into isolate IBCN18, which is virulent towards *Rlm3* and avirulent towards *Rlm4* and *Rlm7* (carrying a functional allele of *AvrLm4-*7) in order to test whether AvrLm4-7 was able to mask Ecp11-1 recognition by Rlm3. Nine IBCN18-*ECP11-1* transformants were obtained. All transformants, as IBCN18, were avirulent on Pixel and virulent on 15.22.4.1 (Fig 6), suggesting that AvrLm4-7 is masking Rlm3*-*mediated recognition of Ecp11-1.

Finally, we complemented one Nz-T4-*ECP11-1* transformant (number 3) with *AvrLm4-7*. Four Nz-T4-*ECP11-1*-*AvrLm4-*7 transformants were obtained. They were avirulent on Pixel, confirming that *AvrLm4-7* was expressed in the transformants. Contrasting with Nz-T4-*ECP11-1* transformants, the Nz-T4-*ECP11-1-AvrLm4-7* transformants were able to cause disease on cultivar 15.22.4.1, confirming that the presence of *AvrLm4-7* is masking Rlm3-mediated recognition of Ecp11-1.

## Discussion

In this study, we determined the crystal structure of *L. maculans* AvrLm5-9 and *F. fulva* Ecp11-1, and obtained a good quality model for AvrLm3 built via the crystal structure of Ecp11-1. Despite their poor sequence similarity, these three effectors are structural analogues of AvrLm4-7. All have a four-stranded β-sheet and helical connections with the same topology. The main differences reside in the conformations of the connections between the strands. Six cysteines involved in disulfide bridges are shared by the three effectors. One disulfide bridge ties together the N- and C-terminal regions, and two others connect the main helical region to the β-sheet. Structure-based pattern searches identified a large number of LARS effector candidates displaying sequence diversity, but likely sharing the same fold. Sequence alignment and 3D model superposition obtained using alphaFold of these candidates shows the strong conservation of six cysteines, which are involved in the aforementioned structure stabilizing disulfide bridges. All of the retrieved sequence analogues are likely compatible with the structures, as confirmed by the structure prediction server I-TASSER. The alignment of the putative analogues highlights a conserved sequence patch, WR(F/L/V)(R/K), with (F/L/V) being a hydrophobic/aromatic residue. These residues are situated at the end of the third β-strand, close to the N and C termini. The tryptophan and arginine are solvent exposed and could provide an interaction surface with plant targets. The hydrophobic (F/L/V) residue is involved in hydrophobic packing of this patch against the β-sheet. Interestingly, site-directed mutagenesis of R^100^ or F^102^ residues resulted in the loss of Rlm7-mediated recognition and abolished the ability of AvrLm4-7 to mask the recognition of AvrLm3 by Rlm3.The latter, however, only occurred in combination with the G^120^R mutation which allowed the effector to escape Rlm4*-*mediated recognition. In contrast to these results, a mutation at residue S^112^ allowed AvrLm4-7 to escape Rlm4 and Rlm7-mediated recognition but also abolished the ability of this effector to mask AvrLm3 from Rlm3*-*mediated recognition. The polymorphic residues identified in AvrLm5-9 and AvrLm3 from *L. maculans* populations and on Ecp11-1 from *F. fulva* isolates are mainly located on the loop regions of the proteins, with the only exceptions being a few amino acid changes (putatively) involved in the switch to virulence towards cultivars with *Rlm3* or *Rlm9* R genes. Remarkably, the positions in the 3D structures of the polymorphic residues G^131^R in AvrLm3 and G^120^R in AvrLm4-7 are very similar, as are I/L^58^H in AvrLm3 and R^55^K in AvrLm5-9, suggesting the same protein regions could be involved in the virulence phenotypes.

Structure-informed pattern searches specifically identified LARS-effectors in phytopathogenic ascomycetes from the Dothideomycetes and Sordariomycetes classes. One or two LARS-effector(s) per species were detected in the Sordariomycetes *Colletotrichum* sp. Two structural analogues were identified in the Dothideomycetes closely related to *L. maculans* ‘brassicae’, specifically *P. tritici repentis* and *P. teres*, while between three and four structural analogues were identified in the species from the complex comprising *L. maculans* ‘brassicae’, suggesting a recent expansion of the LARS family in the species complex. In *L. maculans* ‘brassicae’, thirteen LARS effectors could be detected and their expression during the primary biotrophic stages of oilseed rape cotyledon infection suggests they are *bona fide* effectors. They represent 14 % of the candidate effectors specifically overexpressed during the biotrophic stages of oilseed rape infection (nine LARS effectors among the 63 effector genes in Cluster 2 ‘biotrophy’ defined by Gay et al. [32]). The LARS family also comprises four out of the nine cloned *AVR* genes from *L. maculans*. Most of the *L. maculans* ‘brassicae’ LARS effectors are located in TE-rich regions (9/13) and eight are grouped in three genomic regions as neighbor genes, suggesting their expansion could be partly due to local duplications and that their location in TE-rich compartments could have led to their rapid diversification [34]. Expansions of the LARS family, comprising between 5 to 8 structural analogues, were also detected in two other Dothideomycetes, *M. phaseolina* and *C. cassiicola*. We conclude that LARS effectors probably have a common evolutionary origin and that their expansion in some Dothideomycetes results from duplications, and, at least in the *L. maculans* ‘brassicae’ genome, diversification in TE-rich compartments. Two other structural families of effectors were reported in fungi: the RALPH effectors identified in *Blumeria graminis* and the MAX effectors identified in *Magnaporthe oryzae* (RALPH for RNAse-Like Proteins Associated with Haustoria, [35]; MAX for *Magnaporthe* Avrs and ToxB like, [36]). The RALPH family represents about 25 % of the *B. graminis* predicted effectors and three out of the four AVR effectors identified to date, and most of them are highly expressed during plant infection [10,35,37,38]. Pedersen et al. [35] hypothesized that RALPH effectors originated from an ancestral gene, encoding a secreted ribonuclease, duplicated by TE-driven processes and recently diversified within the grass and cereal powdery mildew lineage. The same way, the MAX family represents between 5 to 10 % of the *M. oryzae* effectors and 25 % of the cloned AVR effectors, and most of them are expressed during early biotrophic stages of rice infection. De Guillen et al. [36] hypothesized that the expansion of the MAX family occurred in a common ancestor of *M. oryzae* and *M. grisea*. The scenario observed for the LARS, RALPH and MAX examples suggests that a wide variety of effectors, without any apparent sequence relationship, could in fact constitute a limited set of structurally conserved effector families and that they have expanded in some fungal lineages or even in several fungal classes.

AvrLm4-7 suppresses Rlm3- and Rlm9-mediated disease resistance. Other cases of effectors with a suppressive function have been described in fungi. In *Fusarium oxysporum* f. sp. *lycopersici*, the Avr1 effector suppresses I-2 and I-3-mediated resistance in tomato triggered respectively by Avr2 and Avr3 [11]. The structure of Avr2 has recently been determined [39], but is unrelated to the structures of AvrLm4-7, AvrLm5-9 or AvrLm3, and the 3D structures of Avr1 and Avr3 are unknown. In the necrotrophic fungus *P. tritici-repentis*, the Host Selective Toxin (HST) ToxA suppresses the activity of other HSTs [40]. Finally, *B. graminis* secretes a suppressor of avirulence (*SvrPm3*) acting on the interaction between the *AVR* gene *AvrPm3* and the barley *R* gene *Pm3* [10]. However, neither in *F. oxysporum*, in *P. tritici-repentis* nor in *B. graminis* have the mechanisms underlying suppressive function yet been determined. In non-fungal models, several mechanisms explaining suppressive interactions have been highlighted. (i) The AVR effector can act downstream of another AVR recognition by an R protein to suppress HR induction: in *Xanthomonas campestris* pv. *vesicatoria*, AvrBsT interacts in the plant cell with SnRK1, an SNF1-related kinase, to inhibit the HR induced by AvrBs1 recognition [41]. (ii) Effectors displaying suppressive interaction can share a common plant target, but differ in their actions on that target: in *Pseudomonas syringae*, the effector proteins AvrRpm1, AvrRpt2 and AvrB target the *Arabidopsis thaliana* protein RIN4, a key regulator of plant immunity [42,43]. While AvrB and AvrRpm1 trigger Rpm1-mediated recognition through phosphorylation of RIN4, AvrRpt2 triggers plant immunity through cleavage of RIN4, thus preventing recognition of AvrB and AvrRpm1 by Rpm1. (iii) Suppressive effectors can directly act on the R proteins: in *Phytophthora infestans*, the IPI-O4 effector suppresses the HR triggered by recognition of IPI-O1 by the potato RB R protein. IPI-O4 interacts with the coiled-coil domain of RB which is also the domain targeted by IPI-O1 [44]. Since AvrLm4-7, AvrLm3 and AvrLm5-9 share the same structural fold, we hypothesize that they could (i) target the same plant components or cellular processes and / or (ii) be recognized by the same R proteins. While we do not have any information on the plant components targeted by AvrLm3, AvrLm5-9 and AvrLm4-7, such a target could be guarded by the R proteins or, in the case of a direct interaction with the R proteins, could be Rlm9 or Rlm3 themselves. *Rlm9*, *Rlm4*, *Rlm7* and *Rlm3* were mapped to the same genetic cluster and could possibly be allelic [45]. *Rlm9* was cloned and found to encode a wall-associated kinase-like (WAKL) protein, a newly described class of Receptor-Like Kinase (RLK) R protein [46]. Using a yeast two-hybrid (Y2H) assay, no direct interaction between the extracellular region of Rlm9 and AvrLm5-9 could be detected. However, a direct interaction between AvrLm5-9 and Rlm9 cannot be excluded since Y2H is not an optimal technique to test interaction with a membrane protein. Haddadi et al. [47] recently cloned *Rlm4* and *Rlm7* and found they corresponded to alleles of *Rlm9*, with the three encoded proteins only differing by a few amino acid residues in the extracellular receptor domain of the WAKL. Larkan et al. [46] had also previously identified at the *Rlm9* locus, in another resistant accession of oilseed rape, a WALK gene that could potentially correspond to *Rlm3*, and being allelic to *Rlm9*. Another example of an allelic R protein that is able to recognize sequence-unrelated AVR effectors with a predicted common fold was recently reported in barley [48], the specificity of recognition being conferred by amino-acid modifications in the LRR domain of the MLA R protein. However, it is currently unknown whether MLA directly interacts with *B. graminis* effectors. Based on the zig-zag model [6], we hypothesize that *Rlm4* and *Rlm7* evolved from *Rlm3* or *Rlm9* in response to the suppressive effect of AvrLm4-7 on AvrLm3 and AvrLm5-9 recognition. We propose a model in which Rlm3 and Rlm9 directly recognize the complex between AvrLm3 (and Ecp11-1) or AvrLm9 and their plant target (or AvrLm3 and AvrLm9 themselves after they have bound to their plant target). AvrLm4-7 would have a higher affinity for the same host target, thereby preventing interaction with AvrLm3 or AvrLm5-9, and thus masking their presence to Rlm3 and Rlm9. Although AvrLm4-7 binds the same host virulence target, we hypothesize that it does not possess the protein region recognized by Rlm3 and Rlm9. Instead, upon binding the plant target, AvrLm4-7 presents a protein region that is recognized by Rlm4 and Rlm7.

We have identified a large structural family of effectors that, in *L. maculans*, are expressed during the early stages of infection and are potentially targeted by R proteins. This structural information on effectors could be used to improve the management and durability of *R* genes in the field. Indeed, among the nine *AVR* genes identified to date in *L. maculans*, four belong to the LARS family. The corresponding *R* genes are, at least in part, present in commercial varieties currently used in the fields (*Rlm7*, *Rlm3*, *Rlm4*, *Rlm9*, *RlmS*). We hypothesize that the presence of *R* genes targeting members of the LARS family potentially exerts a selection pressure on the other members of the family, and that an efficient strategy to improve durability of *R* genes would consist in alternating or pyramiding *R* genes corresponding to different structural classes of effectors. We have also determined that Ecp11-1, a homologue of AvrLm3 and AVR effector candidate from *F. fulva*, is able to trigger *Rlm3*-mediated resistance in oilseed rape. This finding significantly alters our understanding about the degree of host-microbe specificity as developed by Flor in the 1940s [49], in that this is one of the few examples of cross-species effector recognition, as previously mentioned by Stergiopoulos et al. [50] for the recognition of the Avr4 effector from *F. fulva* and of its orthologue in *Mycosphaerella fijiensis* by the Cf4 R protein of tomato. A next step will be to determine whether other homologues identified in Dothideomycetes and Sordariomycetes can also trigger recognition by R proteins of oilseed rape and, in the longer term, to evaluate the possible use of broad-spectrum resistances for multi-pathogen management of diseases.

## Materials and Methods

### *Leptosphaeria maculans*, bacteria and plant growth conditions

The v23.1.3 isolate (also known as JN3) belongs to the race Av1-4-5-6-7-8-10-11-S, i.e. is avirulent towards *Rlm1*, *Rlm4-Rlm8*, *Rlm10-11* and *RlmS*, and is the reference isolate whose genome was sequenced [34] and recently re-assembled and re-annotated [28]. Nz-T4 is a field isolate from New Zealand, IBCN18 is a field isolate from Australia [33], and v45.15 is progeny from a cross between isolates v29.3.1 and 19.2.1 [51]. Nz-T4, IBCN18 and v45.15 were used as recipient isolates for genetic transformation. All fungal cultures were maintained on V8 juice agar medium, and highly sporulating cultures were obtained on V8 juice, as previously described [52].

*E. coli* strain DH5α (Invitrogen) was grown in LB medium at 37 °C. *Agrobacterium tumefaciens* strain C58::pGV2260 was grown in LB medium at 28 °C. Antibiotics were used at the following concentrations: rifampicin 25 μg/ml, ampicillin 50 μg/ml, kanamycin, 100 μg/ml.

*Brassica napus* plants were grown in growth chambers with 16 h of day (22 °C, 80 % humidity) and 8 h of night (18 °C, 100 % humidity).

### Fungal transformation

*A. tumefaciens*–mediated transformation (ATMT) was performed on *L. maculans* as described [53]. Transformants were selected on minimal medium supplemented with 50 μg/ml nourseothricin or 50 μg/ml hygromycin, purified by single conidium isolation and maintained on selective medium.

### Inoculation tests on oilseed rape

All *L. maculans* isolates and transformants were phenotyped for their virulence towards *Rlm3* and *Rlm4* oilseed rape genotypes using a cotyledon-inoculation test [54]. Spore suspensions of each isolate or transformant were inoculated on 10-12 plants of each of the *B. napus* cvs or lines Pixel (*Rlm4*), 15.22.4.1 and 18.22.6.1 (*Rlm3*), Grizzly (*Rlm3*), Columbus (*Rlm1*, *Rlm3*) and 15.23.4.1 (*Rlm7*) [18,22,54,55]. 15.23.4.1 and 18.22.6.1 are two sister lines issued from individual plants from cv. Rangi, that showed a contrasted behavior when inoculated with AvrLm7 or AvrLm3 isolates, respectively. The original lines 23.1.1 and 22.1.1 were generated by three round of selfing of individual plants to reach an homogeneous behaviors of the line [54]. The current lines 15.23.4.1 and 18.22.6.1 results from additional rounds of selfing to multiply and maintain the genotype. Symptoms were scored 12-21 days post-inoculation (DPI) using a semi-quantitative 1 (avirulent) to 6 (virulent) rating scale in which scores 1-3 represent different levels of resistance (from typical HR to delayed resistance) and 4-6 different levels of susceptibility (mainly evaluated by the intensity of sporulation on lesions; [33]). Inoculation tests were repeated twice.

### Bacterial and yeast strains and DNA manipulation

For protein production, all synthetic gene constructs were obtained from Genscript (Piscataway, USA). The DNA sequences coding for AvrLm5-9, AvrLm3 and Ecp11-1 were cloned into a pPICZαA backbone using the *Eco*RI and *Not*I restriction sites in frame with the *Saccharomyces cerevisiae* α–mating factor signal sequence, and under the control of the alcohol oxidase *AOX1* promoter that allows methanol‐inducible expression in *P. pastoris*. *E. coli* strain Top10F’ was used for the amplification of recombinant plasmids. *E. coli* transformants were selected on LBLS plates with 10 mg/ml tetracycline and 25 mg/L Zeocin™. The AvrLm5-9, AvrLm3 and Ecp11-1 constructs were expressed in the eukaryotic expression system *P. pastoris* strain X33 using the EasySelect *Pichia* expression kit (Invitrogen Life Technologies, cat K1740-01) according to the manufacturer’s instructions. These plasmids were used to express AvrLm5-9, AvrLm3 and ECP11-1 with a N-terminal 6His-Trx-tag followed by a TEV proteolytic recognition site to remove the 6His-Trx-tag.

In order to be expressed in *L. maculans*, *ECP11-1* from *F. fulva* was placed under the control of *AvrLm4-7* promoter. The promoter of the *AvrLm4-7* gene was amplified with the primers promAvrLm4-7_Up (GTTTTGGTTAGGTTTAGGGTCT) and promAvrLm4-7_Down BamHI (GAGAGAGGATCCGTTGTTAACTGTCAAAGGGTT). It was then digested with *Nhe*I and *Bam*HI and ligated into a *Spe*I*-Bam*HI-digested pPZPNat1 vector. *ECP11-1* was amplified from its starting codon to its terminator region using primers Ecp11_ATG_EcoU (GAGAGAGAATTCATGTTGTCGTCAGCGAAGACC) and Ecp11_3UTR_XhoL (GAGAGACTCGAGCGACTCCGTAAACTAAGAGATCC) and then digested with *Eco*RI and *Xho*I. This fragment was ligated into the pPZPNat1 vector containing the *AvrLm4-7* promoter and digested with the same enzymes, to obtain the vector pPZPNat1-pAvrLm4-7-Ecp11-1.

A plasmid pPZPNat1 containing *AvrLm4-7* (including its promoter and terminator regions) had been constructed previously [18]. *AvrLm4-7*, from its promoter to its terminator, was transferred into the binary vector pBht2, which carries a hygromycin cassette [56]. pBht2 and pPZPNat1-AvrLm4-7 were digested by *Sac*I and *Sal*I. *AvrLm4-7* was ligated into the pBht2 vector to obtain the vector pBht2-AvrLm4-7.

PCR-mediated site-directed mutagenesis was performed on the plasmid pPZPNat1-AvrLm4-7 that carries v23.1.3 allele of *AvrLm4-7* [18]. Using the pPZPnat1-AvrLm4-7 construct as template, the primers MD1-Up (with a C^299^ instead of G^299^), MD2-Up (with a C^305^ instead of T^305^) or MD3-Up (with a A^336^ instead of C^336^) were used in combination with primer AVR47-JN3-Lo (Table S4) in a PCR reaction using the high fidelity DNA polymerase *Taq* Phusion (Finnzymes) to generate respectively 487-bp fragment MP1, 484-bp fragment MP2 or 452-bp fragment MP3. MP1, MP2 or MP3 was then used as a megaprimer in combination with AVR47-JN3-Up in a *Taq* Phusion PCR reaction to generate 1843-bp products termed v23.1.3-R^100^P, v23.1.3-F^102^S and v23.1.3-S^112^R. Following restriction with *Sna*BI and *Bst*XI and gel purification, the fragments were used to replace the excised corresponding sequence of pPZPNat1-AvrLm4-7. The constructs, termed pPZPNat1-AvrLm4-7-R^100^P, pPZPNat1-AvrLm4-7-F^102^S and pPZPNat1-AvrLm4-7-S^112^R were cloned in *E. coli*, extracted and accuracy of the point mutations checked by sequencing.

### Protein production using *Pichia pastoris* and purification

Proteins were produced in the *P. pastoris* expression system by fed‐batch cultivation. Upon transformation of *P. pastoris* with pPICZαA‐6His-Trx-AvrLm5–9, pPICZαA‐6His-Trx-AvrLm3 and pPICZαA-6His-Trx‐Ecp11-1, transformants were screened for zeocin resistance in shake‐flask cultures. A clone secreting a protein of ~30 kDa (as judged by SDS‐PAGE) was selected for large‐scale production using a fed‐batch mode of cultivation. Recombinant protein was collected from the supernatant, and concentrated. For purification, supernatant was applied onto Ni-NTA resin (Qiagen, Inc.) column affinity chromatography previously equilibrated with Tris–HCl buffer (20 mM Tris–HCl pH 8.0, 300 mM NaCl, 5 % glycerol). Resin was washed with 40 ml of the same buffer and the proteins were eluted using three fractions of 6 ml of the previous buffer supplemented with 100, 200 and 400 mM imidazole. Fractions containing the proteins of interest were pooled and concentrated by ultra-filtration and loaded on Superdex 75 (GE-Healthcare) equilibrated in buffer (20 mM Tris pH 8.0, 300 mM NaCl, 5 % glycerol, 0.5 mM EDTA). The purified fusion proteins were cleaved overnight at 4 °C using the TEV protease. The sample was further purified using an ion exchange column (HiTrap Q HP 5 ml) and the fraction containing the proteins of interest were loaded on a Superdex 75 (GE-Healthcare) gel filtration column equilibrated in buffer (20 mM Tris pH 8.0, 300 mM NaCl, 5 % glycerol). Purified proteins were analyzed by SDS-PAGE (14 %) under denaturing and reducing conditions and protein concentrations were determined with a NanoDrop 2000 spectrophotometer (Thermo Scientific). Finally, fractions containing the proteins of interest were concentrated to 7.2 mg·ml^−1^ or 6 mg·ml^−1^ for AvrLm5-9 and Ecp11-1, respectively. The identity of the recombinant proteins was confirmed by mass spectrometry.

### Crystallization and resolution of the protein structures

Crystals for the AvrLm5-9 protein were obtained at 18 °C from a 0.1 μl:0.2 μl mixture of a 2.7 mg/ml protein solution (stored in 20 mM Tris pH 8.0, 300 mM NaCl and 5 % glycerol) with crystallization liquor composed of 0.7 M sodium acetate, 80 mM nickel sulfate, 0.1 M HEPES pH 7.5. Native crystals were soaked for 10 seconds in 1.2 M sodium acetate, 50 mM nickel sulfate, 0.1 M HEPES pH 7.5 supplemented with 0.2 M sodium iodine and then transferred to a solution composed of 1.25 M sodium acetate, 50 mM nickel sulfate, 0.1 M HEPES pH 7.5 and 30 % of glycerol before flash-cooling in liquid nitrogen. Diffraction data at 2.7 Å resolution were recorded on beam line FIP-BM30A (synchrotron ESRF, France). The space group was P6122 and the asymmetric unit contained one copy of AvrLm5-9 (solvent content of the crystals was 66 %). The structure was determined by the SAD method using the anomalous signal of the iodine atoms and the automated structure solution pipeline implemented in ccp4 with the program CRANK2 [57]. Data were processed using the XDS package [58]. Two iodine sites were located using the program SHELXD [59]. The iterative substructure improvement and phasing of these Iodine sites as well as phasing and hand determination was performed using the REFMAC5, PEAKMAX, MAPRO, Solomon and Multicomb programs. The model was built semi-automatically after density modification using Parrot, REFMAC5 and BUCCANEER [60]. At this step, 123 residues were placed in the electron density with a FOM value of 0.85 and R/R_free_=31/35. The model was further improved by iterative cycles of manual rebuilding using COOT [61] and refinement using REFMAC5 program [62]. Refinement was pursued using 2.14 Å resolution native data recorded on beam line PROXIMA 2 (synchrotron SOLEIL, France). The space group native data (different from those of the I-soaked crystals) was P41212 with one copy of AvrLm5-9 per asymmetric unit. The structure was solved by molecular replacement using the PHASER program [63] and this model was refined using REFMAC5 (R/R_free_ was 22/28 %). The final model contains residues −2 to 121, three nickel atoms, two acetates and 72 water molecules. Statistics for data collection and refinement are summarized in Table S1. The atomic coordinates and structure factors have been deposited into the Brookhaven Protein Data Bank under the accession numbers 7B76 (AvrLm5-9 I-derivative) and 7AD5 (AvrLm5-9 native).

Crystals for the Ecp11-1 protein were obtained at 4 °C from a 0.1 μl:0.2 μl mixture of a 6 mg/ml protein solution (stored in 20 mM Tris pH 8.0, 300 mM NaCl and 5 % glycerol) and crystallization liquor composed of 21 % of PEG550MME, 10 mM zinc sulfate, 0.09 M MES pH 6.5 and 15 % glycerol. Native crystals were flash-cooled in liquid nitrogen and 1.94 Å resolution data were collected on beam line PROXIMA 2 (synchrotron SOLEIL, France). Data were processed using the XDS package [58]. The space group was P2_1_2_1_2_1_ with one Ecp11-1 molecule per asymmetric unit (solvent content of the crystal was 57 %). The structure was determined by the SAD method using the anomalous signal of the zinc atoms and the automated structure solution pipeline implemented in ccp4 with the program CRANK2 [57]. Two Zinc sites were located using the program SHELXD [59]. The iterative substructure improvement and phasing of these zinc sites as well as phasing and hand determination was performed using the REFMAC5, PEAKMAX, MAPRO, Solomon and Multicomb programs. The model was built semi-automatically after density modification using Parrot, REFMAC5 and BUCCANEER [60]. The initial model contained 142 residues with a FOM value of 0.90 and R/R_free_=26/27. The model was further improved by iterative cycles of manual rebuilding using COOT [61] and refinement using REFMAC5 [62]. A native set data at 1.62 Å resolution was recorded on beam line PROXIMA 2 (synchrotron SOLEIL, France) and the model was refined using REFMAC5. This yielded a R/Rfree factor of 19/20 %. Statistics for data collection and refinement are summarized in Table S1. The atomic coordinates and structure factors have been deposited into the Brookhaven Protein Data Bank under the accession numbers 6ZUQ (Ecp11-1 Zn-derivative) and 6ZUS (Ecp11-1 native).

#### Expression analyses using RNAseq data

RNAseq data were previously generated [32]. The expression levels of structural family members were analyzed in *L. maculans* after extraction of RNAseq data corresponding to the following stages of infection: 2, 5, 7, 12 and 15 days after infection of oilseed rape cotyledons (susceptible cultivar Darmor-Bzh). Growth in V8 axenic medium was used as a control. Two biological replicates per condition were tested. We measured the number of RNAseq reads aligned on the different genes after normalization according to the library size (Counts Per Million, CPM). All the genes were detected as being differentially expressed in at least one infection condition compared to the *in vitro* growth condition on V8 solid medium, according to the criterion: Log2 (Fold change)> 2.0 and p-value <0.05.

#### Bio-informatics analyses

For the HMM searches, a homemade database of 163 fungal proteomes was used (the list of the proteomes used can be found in Table S2). The analyses were performed separately for AvrLm4-7, AvrLm5-9 and Ecp11-1. The HMMER search program from the HMMER package v 3.0 was used to perform four iterations with a cut-off E-value of 1 and a minimum overlap of 50 % [30]. The full sequences of the proteins detected by each of these three homology searches were retrieved. Proteins longer than 160 residues were removed. The remaining sequences were aligned with the MAFFT program using parameters --localpair --maxiterate 1000. This final list of sequences was used to build diversity trees using the Neighbor-Joining method using the MEGA6 package. In order to confirm that the retrieved protein analogues have similar structures we used the alphaFold program via the colab server (https://colab.research.google.com/github/sokrypton/ColabFold/blob/main/AlphaFold2.ipynb; [31]) to predict their 3D structure.

Signal peptide predictions were performed using SignalP version 4.1 software [29].

## Acknowledgments

Authors wish to thank all members of the “Effectors and Pathogenesis of *L. maculans*” group. We thank the beamline staff at the ESRF synchrotron (Grenoble France, beamline FIP-BM30A) and Synchrotron SOLEIL (Saclay France, beamline PROXIMA1 and PROXIMA2) for assistance and advice during data collection. David Cobessi (FIP-BM30A beamline) is gratefully acknowledged for his valuable help in the AvrLm5-9 data collection and analysis.

## Supporting information captions

**Fig S1. Purification of the recombinant AvrLm5-9 and Ecp11-1 proteins**.

(A) Size-exclusion chromatogram (Superdex 75 16/60 column; GE Healthcare); elution with 20 mM Tris pH 8.0, 300 mM NaCl, 5 % glycerol. (B) SDS-PAGE gel analysis of the peak fractions of the gel-filtration step. Protein sizes are shown in kDa.

**Fig S2. Diversity of LARS effector structural analogues identified by HMM analyses in *L. maculans* ‘brassicae’ and other phytopathogenic fungi.**

The multiple sequence alignment generated in Figure 5 was used to generate a diversity tree using the Neighbor-joining method. Branch supports are based on 1000 bootstraps and horizontal branch length reflects sequence divergence.

Cg, Colletotrichum gloeosporioides; Ch, Colletotrichum higginsianum; Co, Colletotrichum orbiculare; Cc, Corynespora cassiicola; Ff, Fulvia fulva; Lbb, Leptosphaeria biglobosa ‘brassicae’; Lbt, Leptosphaeria biglobosa ‘thlaspii’; Lmb, Leptosphaeria maculans ‘brassicae’; Lml, Leptosphaeria maculans ‘lepidii’; Mp, Macrophomina phaseolina; Pt, Pyrenophora teres; Ptr, Pyrenophora tritici-repentis; Sl, Stemphylium lycopersici

**Fig S3. 3D models of the LARS effectors identified in *L. maculans***

The 3D structures of the different LARS effectors found by the HMM search were modeled using AlphaFold [31]. Cysteines are presented as sticks and colored in red. No reliable models could be obtained neither for Lmb_jn3_08343 nor for Lmb_jn3_12986.

**Fig S4. Effect of mutations in AvrLm4-7 on the ability to suppress Rlm3-mediated recognition and to induce Rlm4 and Rlm7-mediated recognition**

Wild type isolates G06-E107 (A3a4a7), JN3 (a3A4A7) and JN2 (a3a4A7), as well as G06-E107 transformants carrying AvrLm4-7 alleles with mutations at amino acids R^100^, F^102^, S^112^ and / or G^120^ were inoculated onto cotyledons of cultivars carrying *Rlm3* (15.22.4.1), *Rlm7* (15.23.4.1) or *Rlm4* (Pixel). Pathogenicity was measured 10 and 14 days post-inoculation (DPI). Results are expressed as a mean scoring using the IMASCORE rating comprising six infection classes (IC), where IC1 to IC3 correspond to resistance, and IC4 to IC6 to susceptibility [33]. Error bars indicate the standard deviation of technical replicates.

**Fig S5. Ecp11-1 of *F. fulva* triggers Rlm3-mediated recognition in *L. maculans***

Wild type isolates Nz-T4 (a1a3a4a7), JN3 (A1a3A4A7) and 19.4.24 (A1A3a4a7), as well as Nz-T4 transformants carrying *ECP11-1* or *AvrLm3* were inoculated onto cotyledons of three cultivars carrying *Rlm3* (15.22.4.1, Grizzly and Columbus), *Rlm7* (15.23.4.1) or *Rlm4* (Pixel). 15.23.4.1 is a sister line of 15.22.4.1 issued from individual plants from cv. Rangi, carrying *Rlm7* instead of *Rlm3*. Pathogenicity was measured 15 days post-inoculation. Results are expressed as a mean scoring using the IMASCORE rating comprising six infection classes (IC), where IC1 to IC3 correspond to resistance, and IC4 to IC6 to susceptibility [33]. Error bars indicate the standard deviation of technical replicates.

**Table S1. Data collection and refinement statistics for crystal structures of AvrLm5-9 and Ecp11-1**

**Table S2. Fungal proteome databases used for the HMM analysis**

**Table S3. Characteristics of LARS structural analogues identified in fungi**

**Materials and Methods S1: Selection of efficient protein producer clones of AvrLm5-9 and Ecp11-1**

**Materials and Methods S2: Conditions for native protein production and extraction using *P. pastoris* in fed-batch cultivation**

