## Supplementary material for "A new family of structurally conserved fungal effectors displays epistatic interactions with plant resistance proteins": Supp Table 4

**Table S4: List of PCR primers used for site directed mutagenesis**

|  | **Primers  (5' – 3')** | |
| --- | --- | --- |
| *AVR47-JN3-Lo* |  | TTAAGTGTTGAGTTGCCTAAC |
| *AVR47-JN3-Up* |  | CACTAACCCTAACCTAACCTAT |
| *MD1-Up* |  | cggcgcatagatatccagaattcgttccc |
| *MD2-Up* |  | cgcatagatatcgagaatCcgttcccaaaattc |
| *MD3-Up* |  | ccctatagcagctttagAcagcacctggag |
