## Supplementary Materials and Methods for "A new family of structurally conserved fungal effectors displays epistatic interactions with plant resistance proteins"

### Supplementary experimental procedures

#### Materials and Methods S1: Selection of efficient protein producer clones of AvrLm5-9 and Ecp11-1

The vector pPICZα (Invitrogen) was used to express AvrLm5-9, AvrLm3 or Ecp11-1 in *P. pastoris* without their signal peptide. The constructions contained a histidine tag (6His), Thioredoxin (Trx) and a tobacco etch virus TEV cleavage site at the N terminal part. Preparation of yeast electrocompetent cells and transformation of *P. pastoris* X33 were performed as described by the supplier (Invitrogen, K1710-01). pPICZαA-6His-Trx-Tev-AvrLm5-9, pPICZαA-6His-Trx-Tev-AvrLm3 and pPICZαA-6His-Trx-Tev-Ecp11-1 plasmids were extracted from *E. coli* TOP10F' and digested with *Sac*I. The linear plasmids DNA (5 µg) were transformed into 100 µl of competent X33 cells by pulsed electroporation using a BTX electro cell manipulator (1,500 V, 25 µF, 200 ohms). Transformed cells were incubated on selective YEPD (10 g yeast extract, 20 g peptone, 20 g dextrose and 20 g bacto-agar in 1 liter) agar plates (zeocin™ 100 mg/L) at 30°C for 3-4 days until colonies appeared. Putative multi-copy recombinants were screened by inoculation of 3 µL of each zeocin resistant clones on increasing concentrations of Zeocin™ (500, 1000, and 2000 µg/ml Zeocin™) in the YEPD agar plates. Integration of the gene of interest into the yeast genome was confirmed by direct PCR screening of selected *Pichia* clones in using the 5′AOX1 and 3′ AOX1 primers. Protein expression levels of the selected transformants were explored by cultivation in multiplates-24 wells with 2 mL of buffered methanol complex BMMY medium (10 g yeast extract, 20 g peptone, 100 mM potassium phosphate pH 6, 400 µg biotin, 13.4 g yeast nitrogen base without amino acids and addition of 1%(v/v) methanol 1% twice a day) for 96 hours at 22°C in thermomixed comfort (500 rpm). Protein productions were estimated by SDS PAGE analysis of the filtrated and concentrated supernatants (cut off 10 kDa).

**Materials and Methods S2: Conditions for native protein production and extraction using *P. pastoris* in fed-batch cultivation**

The best secreting clones were cultivated in high cell density using a fed-batch mode of cultivation. Large-scale cultures were done in a bioreactor (DASGIP) in presence of 1 Liter of the modified buffered glycerol complex BMGY medium, containing all the components as BMGY medium except methanol is replace by 4 % (v/v) of glycerol. Ammonia 15 % (v/v) was used both to maintain the pH at 6 and as source of nitrogen. The temperature was set at 28°C, oxygenation was 1 volume of sterile air per volume of medium per minute and oxygen saturation was maintained over 10% by vigorous stirring (1200 rpm). Upon depletion of glycerol, a pulse with an identical concentrated medium was given. When the entire carbon source was consumed, induction by methanol was initiated with a gradual supply of the feed-medium. The synthetic feed-medium used for fed-batch culture contained 780 g methanol, trace elements 5X PTM1 [$H_3BO_3$ 0.1 mg, $CuSO_4.5H_2O$, 30 mg; KI 0.5 mg, $MnSO_4.H_2O$, 15 mg, $Na_2MoO_4$, 1 mg, $ZnCl_2$, 100 mg, $FeSO_4.7H_2O$ 325 mg, $H_2SO_4$ (96%) 0.1 mL and biotin 1 mg per liter]. The feeding profile was monitored using a computer-controlled pump to maintain the biomass yield $Y_{X/S}$ of 0.3 g biomass per gram of methanol and a theoretical specific growth rate of about 0.015 per hour. Induction went on for three days (1.5 generation cells) after the first addition of methanol. Biomass was determined by measurement of optical density at 600 nm and dry cell weight of yeast suspensions (according to a correspondence of 0.48 g/L dry cell weight per $DO_{600nm}$ unit).

Supernatants of the cultures were collected by centrifugation at 5000 rpm for 20 min and then filtrated (0.5 µM) and diafiltrated (10 kDa) against appropriated buffer (20 mM Tris pH 8, 300 mM NaCl, 5% v/v glycerol) by tangential flow ultrafiltration on hollow fiber cartridges (GE Quixstand system). The supernatant was used to purify the recombinant proteins.
