## Supplementary material for "A new family of structurally conserved fungal effectors displays epistatic interactions with plant resistance proteins": Supp Figure 1

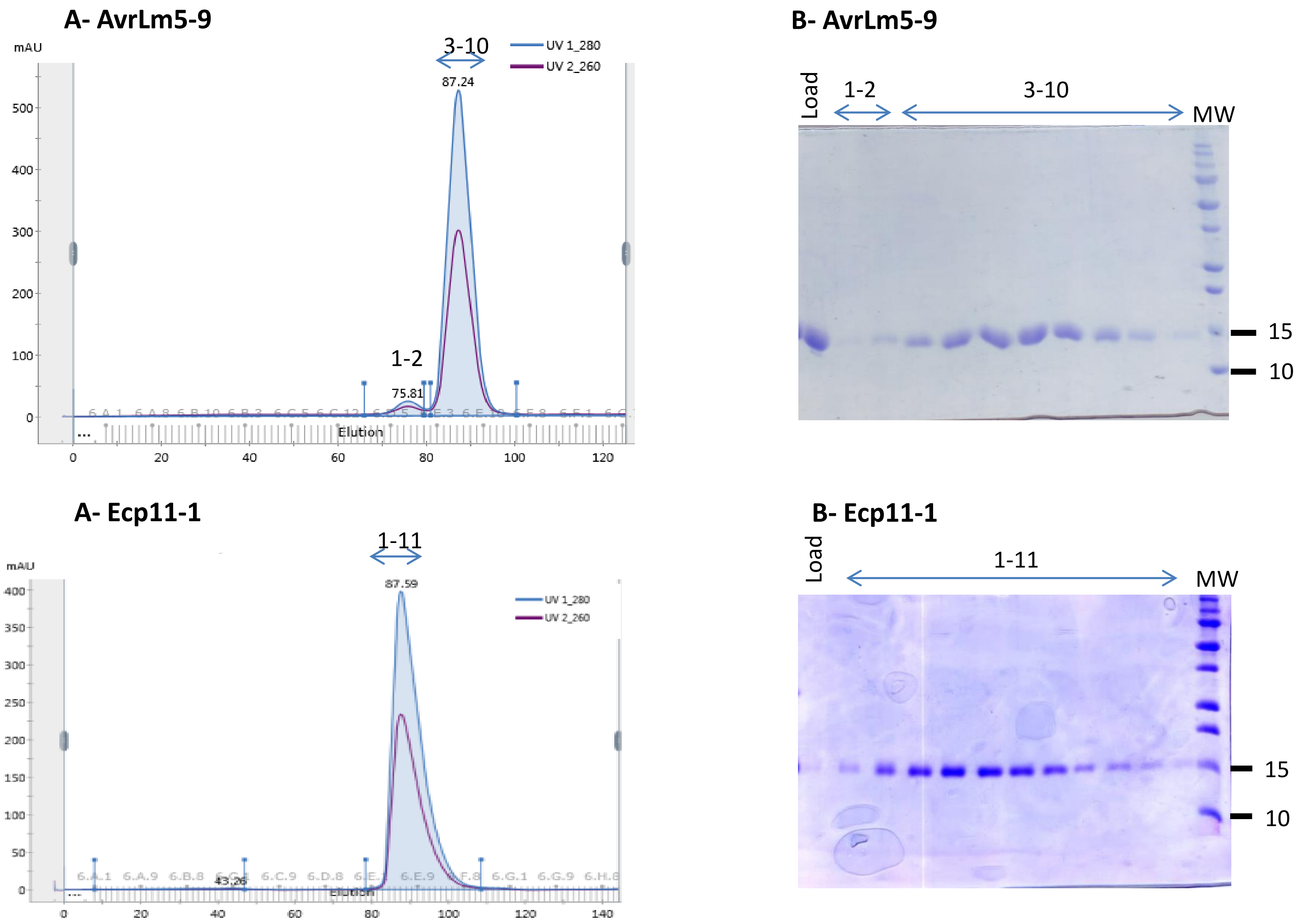


**Fig S1. Purification of the recombinant AvrLm5-9 and Ecp11-1 proteins.**
(A) Size-exclusion chromatogram (Superdex 75 16/60 column; GE Healthcare); elution with 20 mM Tris pH 8.0, 300 mM NaCl, 5 % glycerol. (B) SDS-PAGE gel analysis of the peak fractions of the gel-filtration step. Protein sizes are shown in kDa.
