## Supplementary material for "A new family of structurally conserved fungal effectors displays epistatic interactions with plant resistance proteins": Supp Figure 2

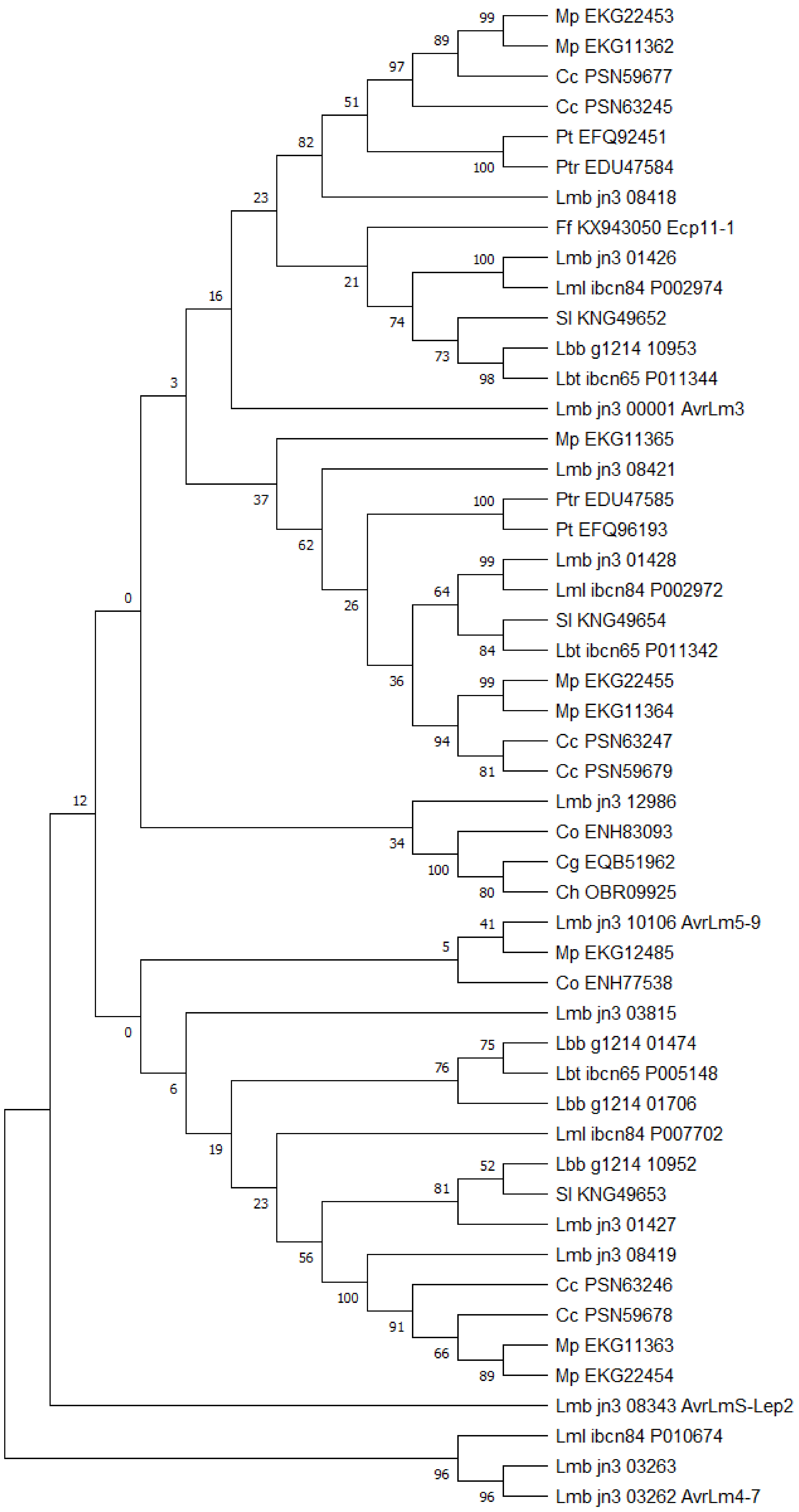


**Fig S2. Diversity of LARS effector structural analogues identified by HMM analyses in *L. maculans* 'brassicae' and other phytopathogenic fungi.**
The multiple sequence alignment generated in Figure 5 was used to generate a diversity tree using the Neighbor-joining method. Branch supports are based on 1000 bootstraps and horizontal branch length reflects sequence divergence.
Cg, *Colletotrichum gloeosporioides*; Ch, *Colletotrichum higginsianum*; Co, *Colletotrichum orbiculare;* Cc, *Corynespora cassiicola*; Ff, *Fulvia fulva*; Lbb, *Leptosphaeria biglobosa* 'brassicae'; Lbt, *Leptosphaeria biglobosa* 'thlaspii'; Lmb, *Leptosphaeria maculans* 'brassicae'; Lml, *Leptosphaeria maculans* 'lepidii'; Mp, *Macrophomina phaseolina*; Pt, *Pyrenophora teres;* Ptr, *Pyrenophora tritici-repentis*; Sl, *Stemphylium lycopersici*
