## Supplementary material for "A new family of structurally conserved fungal effectors displays epistatic interactions with plant resistance proteins": Supp Figure 3

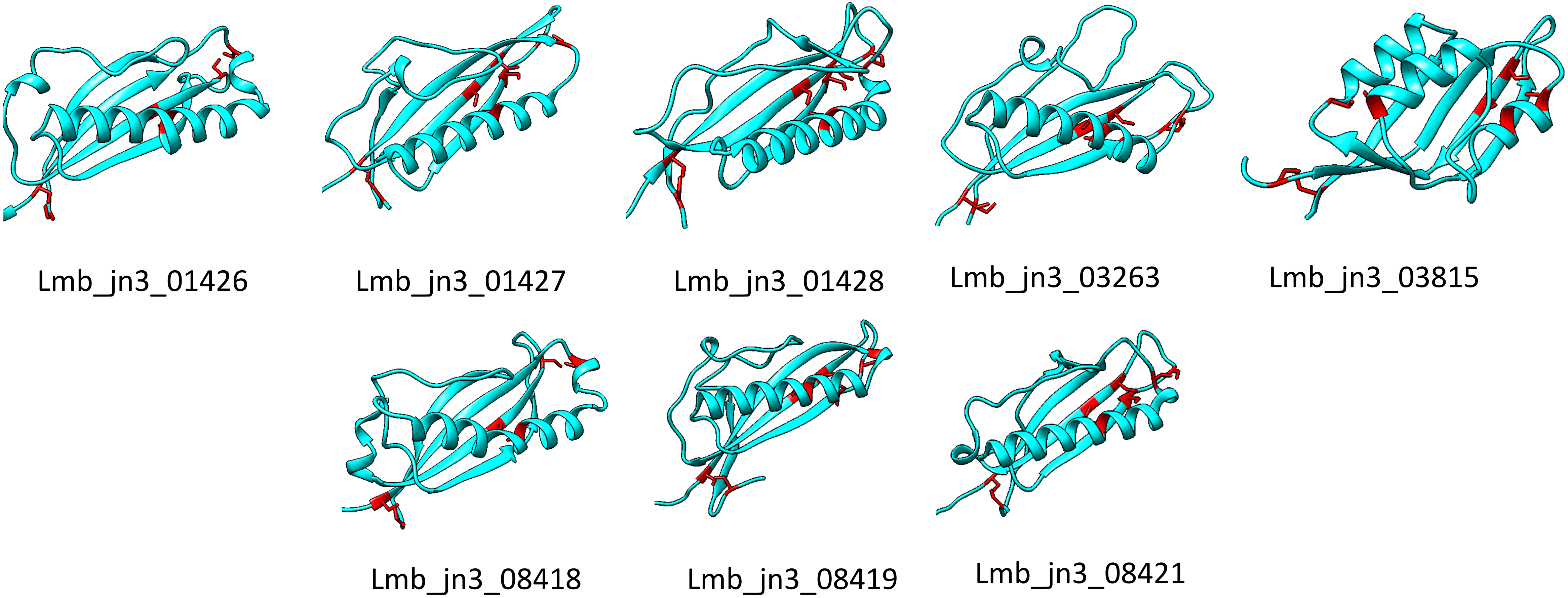


**Fig S3. 3D models of the LARS effectors identified in *L. maculans***

The 3D structures of the different LARS effectors found by the HMM search were modeled using AlphaFold (31). Cysteines are presented as sticks and colored in red. No reliable models could be obtained neither for Lmb_jn3_08343 nor for Lmb_jn3_12986
