## Supplementary material for "A new family of structurally conserved fungal effectors displays epistatic interactions with plant resistance proteins": Supp Figure 4

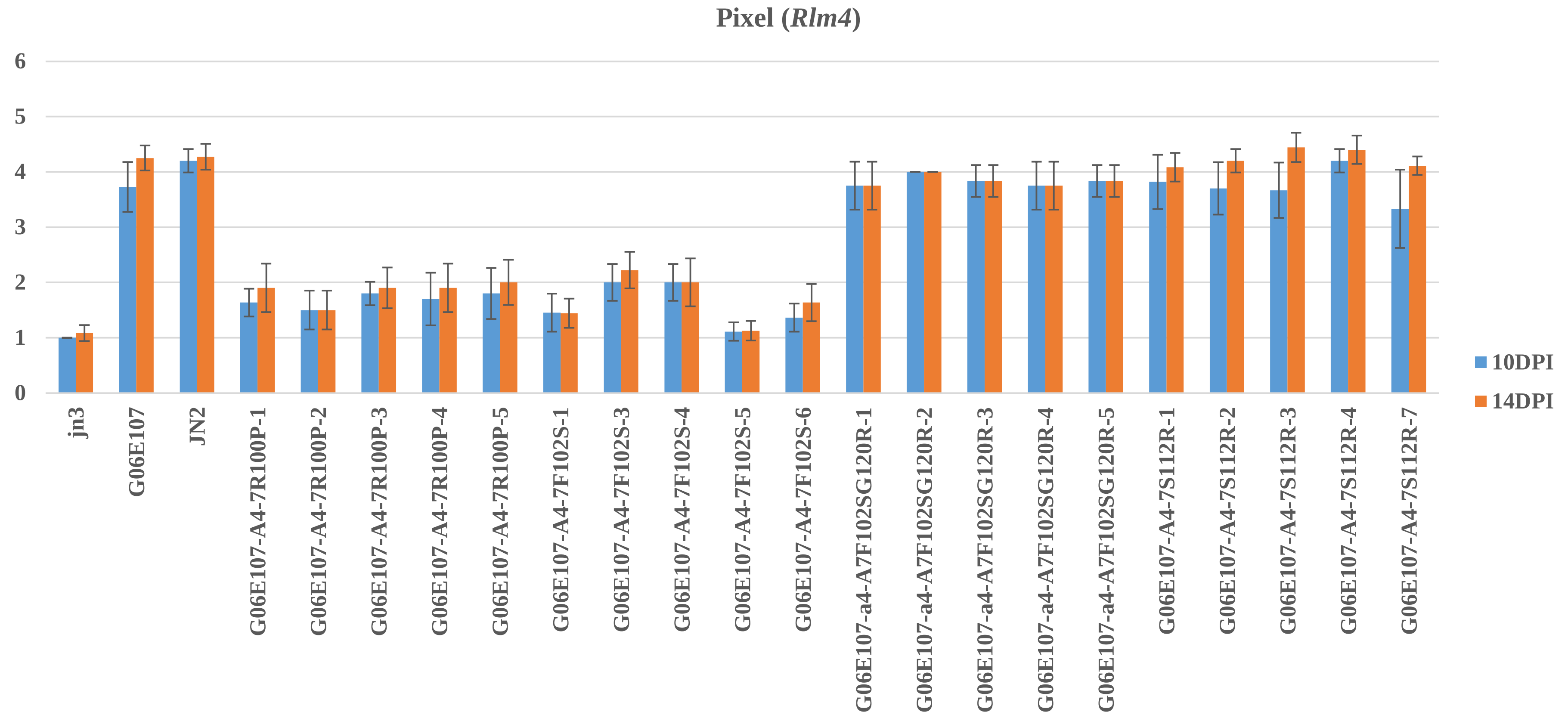


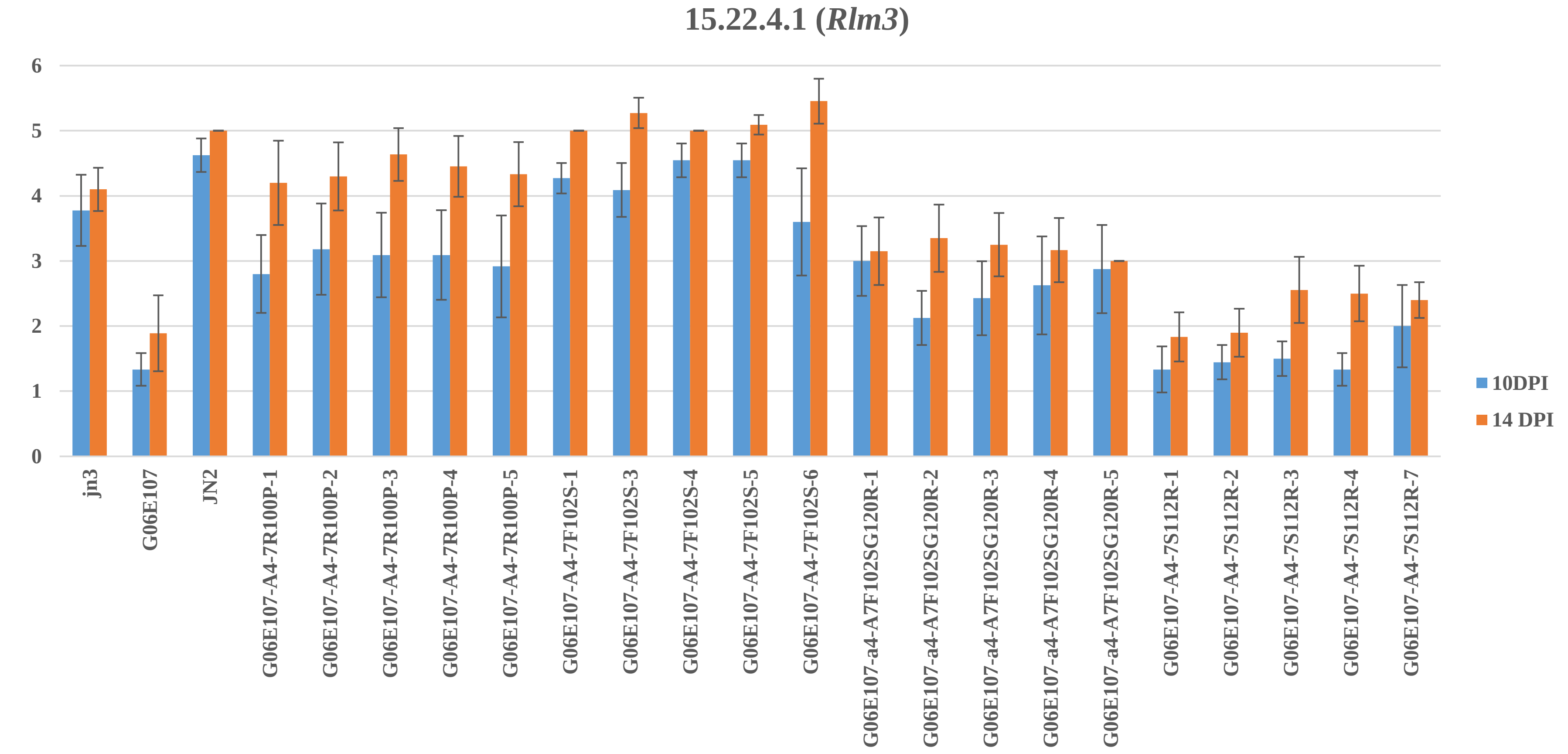


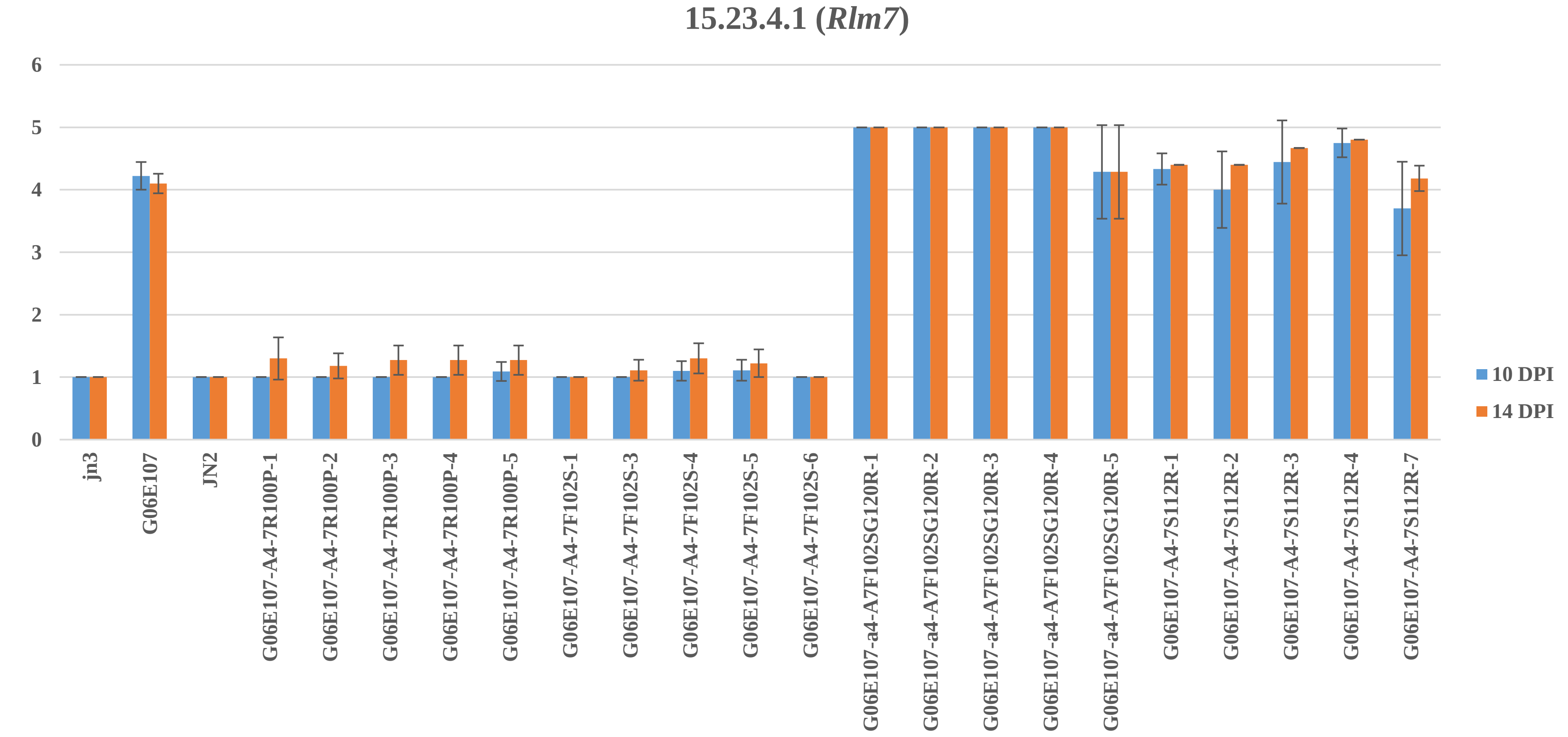


**Fig S4. Effect of mutations in AvrLm4-7 on the ability to suppress Rlm3-mediated recognition and to induce Rlm4 and Rlm7-mediated recognition**

Wild type isolates G06-E107 (A3a4a7), JN3 (a3A4A7) and JN2 (a3a4A7), as well as G06-E107 transformants carrying AvrLm4-7 alleles with mutations at amino acids $R^{100}$, $F^{102}$, $S^{112}$ and / or $G^{120}$ were inoculated onto cotyledons of cultivars carrying *Rlm3* (15.22.4.1), *Rlm7* (15.23.4.1) or *Rlm4* (Pixel). Pathogenicity was measured 10 and 14 days post-inoculation (DPI). Results are expressed as a mean scoring using the IMASCORE rating comprising six infection classes (IC), where IC1 to IC3 correspond to resistance, and IC4 to IC6 to susceptibility [33]. Error bars indicate the standard deviation of technical replicates.
