## Supplementary material for "A new family of structurally conserved fungal effectors displays epistatic interactions with plant resistance proteins": Supp Figure 5

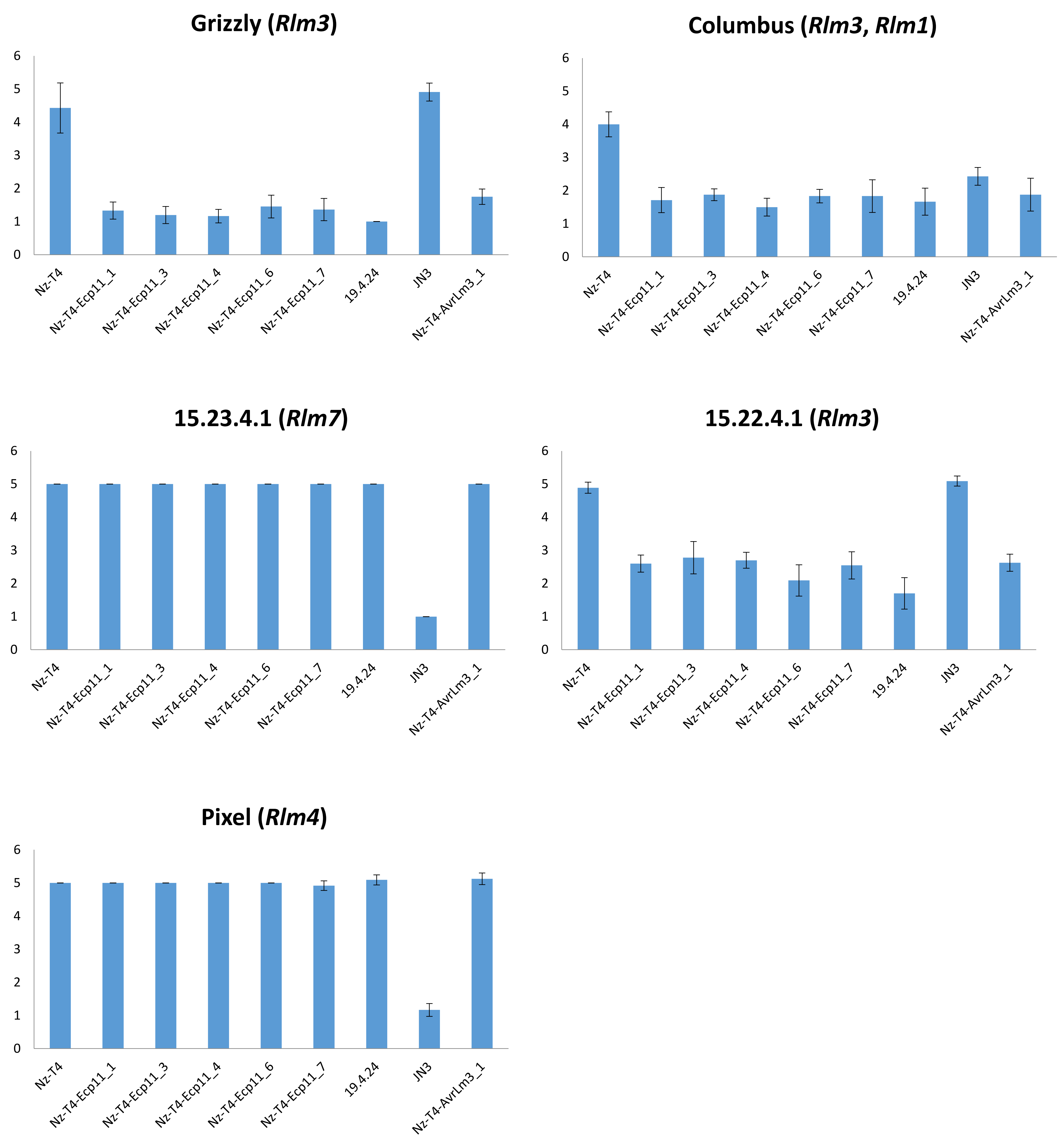


**Fig S5. Ecp11-1 of *F. fulva* triggers Rlm3-mediated recognition in *L. maculans***

Wild type isolates Nz-T4 (a1a3a4a7), JN3 (A1a3A4A7) and 19.4.24 (A1A3a4a7), as well as Nz-T4 transformants carrying *ECP11-1* or *AvrLm3* were inoculated onto cotyledons of three cultivars carrying *Rlm3* (15.22.4.1, Grizzly and Columbus), *Rlm7* (15.23.4.1) or *Rlm4* (Pixel). 15.23.4.1 is a sister line of 15.22.4.1 issued from individual plants from cv. Rangi, carrying *Rlm7* instead of *Rlm3*. Pathogenicity was measured 15 days post-inoculation. Results are expressed as a mean scoring using the IMASCORE rating comprising six infection classes (IC), where IC1 to IC3 correspond to resistance, and IC4 to IC6 to susceptibility [33]. Error bars indicate the standard deviation of technical replicates.
