## Supplementary material for "A new family of structurally conserved fungal effectors displays epistatic interactions with plant resistance proteins": Supp Table 1

**Table S1*.* : Data collection and refinement statistics for crystal structures of AvrLm5-9 and Ecp11-1**

***Crystallographic data collection***

|  | AvrLm5-9  (Iodide derivative) | AvrLm5-9 | Ecp11-1  (zinc derivative) | Ecp11-1 |
| --- | --- | --- | --- | --- |
| X-ray source | FIP-BM30A | PROXIMA2 | PROXIMA2 | PROXIMA2 |
| Wavelength (Å) | 1.4878 | 0.9801 | 1.2819 | 0.9734 |
| Unit-cell parameters (Å, °) | ***a*** = ***b*** = 57.9,  ***c*** = 212.81  ***α*** = ***β*** =90, ***γ*** = 120 | ***a*** = ***b*** = 57.24,  ***c*** = 119.53  ***α*** = ***β*** = ***γ*** = 90 | ***a*** = 40.76, ***b*** = 53.63, ***c*** = 86.27  ***α*** = ***β*** = ***γ*** = 90 | ***a*** = 40.64, ***b*** = 53.64, ***c*** = 86.15  ***α*** = ***β*** = ***γ*** = 90 |
| Space group | P6_1_22 | P4_1_2_1_2 | P2_1_2_1_2_1_ | P2_1_2_1_2_1_ |
| Resolution limits^†^ (Å) | 48.81 – 2.7  (2.86 -2.7) | 41.34 - 2.14  (2.27 - 2.14) | 43.14 - 1.94  (2.06 – 1.94) | 45.55 - 1.62  (1.72-1.62) |
| Number of observations^†^ | 123040 (19352) | 143964 (22293) | 184783 (27217) | 325275 (50766) |
| Number of unique reflections | 10992 (1785) | 11526 (1742) | 26981 (4248) | 46421 (7420) |
| R-meas^†^ (%) | 29.9 (222.1) | 23.1 (184.1) | 12.1 (143.5) | 10.2 (141.7) |
| Completeness^†^ (%) | 99.8 (99.0) | 99.4 (96.3) | 99.5 (97.1) | 99.7 (98.5) |
| I/σ† (I) | 8.8 (1.2) | 8.6 (1.1) | 10.24 (1.13) | 10.56 (0.86) |
| CC (1/2) | 99.2 (65.4) | 99.7 (81.7) | 99.7 (41.3) | 99.8 (60.4) |
| SigAno^++^ | 0.95 (1.18) | - | 1.15 (1.02) | - |
| Anomal Corr^++^ | 23 (45) | - | 39 (30) |  |

^†^ Values in parentheses refer to the highest resolution shell.

^††^ Values in parentheses refer to 3.62 Å and 2.2 Å resolution for AvrLm5-9 and Ecp11-1, respectively.

***Refinement***

|  | AvrLm5-9 | Ecp11-1 |
| --- | --- | --- |
| Number of non-hydrogen atoms (Protein/other/water) | 993/11/70 | 1176/35/159 |
| R/R_free_ (%) | 22.4/27.5 | 18.5/19.7 |
| R.M.S.D. Bonds (Å)/angles (°) | 0.008/1.56 | 0.017/2.06 |
| Average temperature factors (Protein/other) | 53.9/62.2 | 27.6/42.3 |
